# Blood metabolome signature predicts gut microbiome α-diversity in health and disease

**DOI:** 10.1101/561209

**Authors:** Tomasz Wilmanski, Noa Rappaport, John C. Earls, Andrew T. Magis, Ohad Manor, Jennifer Lovejoy, Gilbert S. Omenn, Leroy Hood, Sean M. Gibbons, Nathan D. Price

## Abstract

Defining a ‘healthy’ gut microbiome has been a challenge in the absence of detailed information on both host health and microbiome composition. Here, we analyzed a multi-omics dataset from hundreds of individuals (discovery n=399, validation n=540) enrolled in a consumer scientific wellness program to identify robust associations between host physiology and gut microbiome structure. We attempted to predict gut microbiome α-diversity from nearly 1000 analytes measured from blood, including clinical laboratory tests, proteomics and metabolomics. While a broad panel of 77 standard clinical laboratory tests and a set of 263 proteins from blood could not accurately predict gut microbial α-diversity, we found that 45% of the variance in microbial community diversity was explained by a subset of 40 blood metabolites, many of microbial origin. This relationship between the host metabolome and gut microbiome α-diversity was very robust, persisting across disease conditions and antibiotics use. Several of these novel metabolic biomarkers of gut microbial diversity were previously associated with host health (e.g. cardiovascular disease risk, diabetes, and kidney function). A subset of 11 metabolites classified participants with potentially problematic low α-diversity (ROC AUC=0.88, Precision-Recall AUC=0.76). Relationships between host metabolites and α-diversity remained consistent across most of the Body Mass Index (BMI) spectrum, but were modified in extreme obesity (class II/III, but not class I), suggesting a significant metabolic shift. Out-of-sample prediction accuracy of α-diversity from the 40 identified blood metabolites in a validation cohort, whose microbiome samples were analyzed by a different vendor, confirmed the robust correspondence between gut microbiome structure and host physiology. Collectively, our results reveal a strong coupling between the human blood metabolome and gut microbial diversity, with implications for human health.

## Background

Changes in the composition or structure of the gut microbiome have been associated with an increasing number of human diseases, including diabetes, colorectal cancer and complex gastrointestinal disorders such as inflammatory bowel disease (IBD) ^1,2^. Each diverse bacterial community residing in each human gut can play various roles in human health, from metabolizing nutrients and bile acids to secreting hormones and maintaining healthy immune function ^3–5^. A recent study demonstrated that the fecal metabolome can account for close to 68% of the variance in inter-individual gut microbiome composition, demonstrating the strong association between the metabolic output and gut microbial structure ^6^. While fecal metabolites might be more reflective of the direct metabolic output of the microbiota, blood metabolites provide a window into which of these compounds make it into circulation to potentially impact host metabolism and health. The emergence of untargeted metabolomics has increased our understanding of the blood metabolome and has led to identification of unique molecules in circulation that are generated by the gut microbiota and that may exert biological effects in the host ^7,8^. However, many questions remain about how closely the blood metabolome reflects the composition of the gut microbiome, what metabolites in particular are most strongly associated with gut microbial diversity, and under what physiological conditions the gut microbiome-host metabolome relationship is perturbed.

Shannon diversity is a common metric of α-diversity (within-sample diversity) that summarizes taxonomic richness (how many species are represented) and evenness (the degree to which these species are represented at the same level) (see Methods), and has been suggested as a marker for microbiome health. Lower Shannon diversity, relative to healthy controls, has been reported in individuals diagnosed with IBD, as well as enteric infections caused by *Clostridium difficile* or other bacterial pathogens ^1,9,10^. Conversely, increased Shannon diversity has been reported in hunter-gatherer communities compared to individuals in industrialized countries, suggesting an unprocessed diet, an active lifestyle, or increased exposure to intestinal parasites may contribute to a more diverse gut microbiome ^11,12^. Bristol Stool Scores indicative of constipation (i.e. harder stool) have also been associated with higher Shannon diversity ^13^, which suggests higher diversity may not be optimal for human health above some threshold. Therefore, the debate on how to define an optimal gut α-diversity, and whether such an optimum exists, continues.

Regular, efficient monitoring of gut α-diversity in a clinical setting could have important implications for diagnosing, monitoring and treating disease. Such a test could be used to assess recovery from gastrointestinal illness, monitor gut health in chronic conditions such as IBD, or optimize interventions for improving gastrointestinal health. Commonly used medications such as antibiotics and proton pump inhibitors have been shown to decrease gut α-diversity and increase the risk of enteric infections, including recurrent *Clostridium difficile* infection (rCDI) ^14–16^. rCDI kills thousands of patients each year ^17^ and is characterized by a substantial decrease in Shannon diversity relative to patients experiencing an initial CDI ^10^. Fecal microbiota transplants (FMTs) are the most promising line of treatment for rCDI^18^, however, presently they are only approved for cases with multiple CDI recurrences and where other treatment options have fallen short. By creating an easy, reliable method for monitoring gut α-diversity, patients at high risk for rCDI could be identified earlier and FMT administered as first line of treatment, resulting in improved patient outcomes.

Perhaps nowhere is the controversy about the gut microbiome and its relation to health greater than for obesity ^19^. Obesity is an established risk factor for many diseases, including cardiovascular disease, diabetes, and several types of cancers ^20^. Changes in gut microbiota composition, including shifts in α-diversity, have been linked to obesity, although these associations are inconsistent across studies ^1,19^. Chronic inflammation underlies many obesity-related health risks and is believed to be mediated by cytokines produced in the adipose tissue and/or by a low-fiber diet that promotes a pro-inflammatory microbiota ^21^. Even in the absence of metabolic abnormalities, obesity is associated with higher risk for cardiovascular and autoimmune diseases ^22,23^, prompting a deeper investigation into the host-microbiome relationship across the BMI spectrum.

Previously, our research group published analyses incorporating proteomic, metabolomic, genetic, and microbiome data on 108 participants in the context of health and wellness ^24^. Through a partnership between the Institute for Systems Biology (ISB) and a spin-out company, Arivale, we have greatly expanded the cohort through individuals who have consented to allow their de-identified data to be used for scientific discovery. This wealth of data provides a unique research opportunity for investigating the host-microbiome interface, revealing novel insight into how the gut microbiota corresponds to human health. In this study, we utilized dense phenotyping of Arivale participants to assess the relationship between a multitude of blood analytes and gut microbial Shannon diversity. We demonstrate that a relatively small subset of 40 of the 659 plasma metabolites measured can be used to predict Shannon diversity in the gut, which indicates a strong association between host physiology and the structure of the gut microbiome. We find little-to-no correspondence between 77 clinical lab analytes and Shannon diversity, which suggests that biomarkers that are relevant to the state of the microbiome are not routinely measured in clinical settings. Our results further indicate that specific metabolite-Shannon diversity associations vary across BMI classes, particularly under severe obesity. A great majority of the identified observations were confirmed in a validation cohort, that consists of participants who joined the Arivale program at an earlier date and whose stool and blood samples were analyzed by a different vendor, demonstrating the robustness of the identified relationships despite the added variability seen across different data generating pipelines ^25^. Collectively, the present study provides novel insight into the complex interactions between host health and the gut microbiome, with particular focus on the blood metabolome.

## Methods

### Cohort

Subscribers to the Arivale Scientific Wellness^™^ program (Arivale, Inc., Seattle, WA) were included in the study if they gave permission to use their de-identified data for scientific discovery and met the inclusion criteria. The inclusion criteria required participants to have their initial blood draw taken within 21 days of their stool sample (no. of days from blood draw median=0, interquartile range (IQR)= −5 to 6). This cutoff was chosen to prevent large gaps between when samples were collected for blood metabolomics and gut microbiome analysis. To be included in the discovery cohort, participants were further required to have their gut microbiome analyzed by DNA Genotek, one of two microbiome vendors used in the program, and their blood clinical labs by Labcorp of America (LCA) (N=399). Demographic information on the cohort is provided in Table 1, where participants were stratified based on BMI into four classes: normal weight (18<BMI<25), overweight (25≤BMI<30), World Health Organization (WHO) obese category I (obese I) (30≤BMI<35) and WHO obese categories II and III (35≤BMI). Our cohort was predominantly white with a mean age of 47 years and a greater proportion of females (72%) than males. Overall our cohort represents a broad range of adults, with markers of health decreasing with increasing BMI. An additional 540 participants who met the 21-day cut off criteria, but whose gut microbiome was analyzed by Second Genome were included as a validation cohort. Demographic information on the validation cohort is provided in Supplementary Table 1.

**Table 1.** Baseline discovery cohort characteristics overall and by BMI class. Values with different superscript letters are significantly different (P<0.05).

|  | Normal Weight<br>(n=142) | Overweight<br>(n=134) | Obese I<br>(n=66) | Obese II/III<br>(n=57) | Total<br>(N=399) |
| --- | --- | --- | --- | --- | --- |
| **Mean Age**<br>**(SD)** | 45.0<sup>a</sup><br>(9.1) | 47.7<sup>b</sup><br>(10.6) | 48.6<sup>b</sup><br>(10.6) | 48.8<sup>b</sup><br>(9.7) | 47.0<br>(10.1) |
| **Male**<br>**(%)** | 27.5<sup>a</sup> | 37.3<sup>b</sup> | 27.3<sup>a</sup> | 7.0<sup>c</sup> | 27.8 |
| **Non-white**<br>**(%)** | 26.0 | 24.6 | 21.2 | 24.6 | 24.6 |
| **Median BMI**<br>**(IQR)** | 22.8<sup>a</sup><br>(21.1-24.1) | 27.2<sup>b</sup><br>(26.2-28.5) | 31.9<sup>c</sup><br>(30.7-33.2) | 40.9<sup>d</sup><br>(36.9-45.8) | 26.8<br>(23.0-29.1) |
| **Mean HDL (mg/dL)**<br>**(SD)** | 67.7<sup>a</sup><br>(17.8) | 60.8<sup>b</sup><br>(15.9) | 55.5<sup>c</sup><br>(12.2) | 52.7<sup>c</sup><br>(13.9) | 61.2<br>(16.7) |
| **Mean Blood**<br>**Triglycerides**<br>**(mg/dL) (SD)** | 81.5<sup>a</sup><br>(32.9) | 96.6<sup>b</sup><br>(39.0) | 111.0<sup>c</sup><br>(41.9) | 131.3<sup>d</sup><br>(65.8) | 98.6<br>(45.7) |
| **Median IL-6**<br>**(pg/mL) (IQR)** | 0.70<sup>a</sup><br>(0.70-1.4) | 0.95<sup>b</sup><br>(0.70-1.7) | 1.5<sup>c</sup><br>(0.80-2.3) | 3.1<sup>d</sup><br>(2.0-5.2) | 1.1<br>(0.70-2.3) |
| **Median**<br>**C-Reactive Protein**<br>**(ng/mL)(IQR)** | 0.75<sup>a</sup><br>(0.39-1.12) | 1.12<sup>b</sup><br>(0.55-2.88) | 2.20<sup>c</sup><br>(1.10-3.16) | 4.50<sup>d</sup><br>(2.53-8.82) | 1.27<br>(0.62-2.95) |
| **Self-reported**<br>**Probiotic Use (%)** | 14.1 | 11.9 | 13.6 | 8.8 | 12.5 |
| **Mean Shannon**<br>**Diversity (SD)** | 4.11<sup>a</sup><br>(0.40) | 4.02<sup>b</sup><br>(0.41) | 3.89<sup>b,c</sup><br>(0.50) | 3.77<sup>c</sup><br>(0.50) | 4.00<br>(0.45) |
| **Mean PD whole tree**<br>**Diversity (SD)** | 35.7<sup>a</sup><br>(6.33) | 34.1<sup>b</sup><br>(6.21) | 32.7<sup>b</sup><br>(7.64) | 30.0<sup>c</sup><br>(6.78) | 33.86<br>(6.86) |
| **Mean Chao1**<br>**(SD)** | 971.8<sup>a</sup><br>(180.2) | 921.9<sup>b</sup><br>(185.0) | 898.2<sup>a,b</sup><br>(229.0) | 817.0<sup>c</sup><br>(218.5) | 920.7<br>(202.7) |

### Clinical Laboratory Tests

All blood draws for all assays were performed by trained phlebotomists at LCA or Quest service centers. Blood samples for clinical labs were obtained at the same time as the metabolomics blood draw, within 21 days of the gut microbiome sample. Participants were required to avoid alcohol, vigorous exercise, aspartame, and monosodium glutamate 24 hours prior to the blood draw, and to begin fasting 12 hours in advance. All clinical labs were analyzed by LCA in the discovery cohort, and by Quest (62% of samples) or LCA (38%) in the validation cohort. At the time of each blood draw, weight and height were measured, and BMI was calculated using the formula: 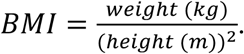 A threshold of less than 5% missing values was set for each clinical lab, which was passed by 77 different analytes. Median imputation was performed on the resulting analytes, which were then used for further analysis.

### Proteomics

Plasma protein levels were measured using the ProSeek Cardiovascular II, Cardiovascular III, and Inflammation arrays (Olink Biosciences, Uppsala, Sweden), processed and batch corrected as described before ^26^. For analysis, a threshold of less than 5% missing values was set for each protein, which was passed by 263 different analytes. The number of participants who met the initial inclusion criteria and whose proteomics measurement was obtained within 21 days of the gut microbiome sample was 262. Missing values for the proteins were imputed to be the minimum observed value for that protein.

### Metabolomics

Metabolon (North Carolina) conducted the metabolomics assays on participant plasma samples. Sample handling, quality control, and data extraction, along with biochemical identification, data curation, quantification, and data normalizations have been previously described ^27^. For analysis, the raw metabolomics data were median scaled within each batch such that the median value for each metabolite was one. To adjust for possible batch effect, further normalization across batches was performed by dividing the median-scaled value of each metabolite by the corresponding average value for the same metabolite in QC samples of the same batch. Missing values for metabolites were imputed to be the minimum observed value for that metabolite. Values for each metabolite were log transformed. A total of 659 different plasma metabolites were measured for each participant in the discovery and validation cohorts.

### Microbiome

Stool specimens were taken at participants’ homes using a standardized kit supplied by DNAgenotek (Ottawa, ON, Canada) in the discovery cohort and Second Genome (South San Francisco, CA) in the validation cohort. Microbial DNA was isolated from 250 mL of homogenized stool, using an automated protocol and MoBio’s PowerMag (+ClearMag) microbiome DNA isolation kit on the KingFisher Flex instrument. The extraction protocol involved a precursory bead-beating step with glass beads and plate shaker for recovery of more DNA from a more diverse microbial community. Concentrations of extracted DNA from each sample were determined by Qubit measurement, and an estimate of sample purity was determined with spectrophotometry by measuring the A260/A280 absorbance ratio. Gut microbiome sequencing data in the form of FASTQ files were provided based on either the 300-bp paired-end MiSeq profiling of the 16S V3+V4 region (DNAgenotek) or 250-bp paired-end MiSeq profiling of the 16S V4 region (Second Genome). Operational taxonomic unit (OTU) read counts were calculated using the QIIME pipeline ^28^ (version 1.9.1; default parameters) with closed-reference OTU picking against the Greengenes database (version 13_08) ^29^. Rare OTUs, defined here as those not representing 0.01% of at least one sample, were removed. Remaining OTU counts were unit normalized. For α-diversity calculations, OTUs were rarefied at 30,000 reads. Gut α-diversity was calculated by 1) Shannon’s index, calculated by 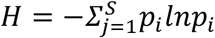, where *p_i_* is the proportion of the community represented by OTU_i_, 2) Chao1, calculated by 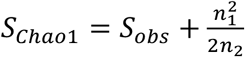, where *S*_obs_ is the number of observed species, n_1_ is the number of singletons (species captured once), and n_2_ is the number of doubletons (species captured twice) ^30^, and by 3) PD whole tree - defined as the minimum total length of all the phylogenetic branches required to span a given set of taxa on the phylogenetic tree ^31^. Participants with Shannon diversity values greater or less than 3.0 standard deviations from the mean were removed prior to analysis (n=4 for the discovery cohort, n=6 for the validation cohort). For microbiome genera-metabolite correlation analysis, only genera that had less than 5% zero values and a mean greater than 5 counts were used. This decreased the number of genera from 550 to 96. Spearman correlations were then used to compare each of the 11 metabolites retained by all 10 LASSO models with each of the 96 genera, correcting for multiple hypothesis testing (FDR<0.05).

### Gastrointestinal Health Metrics

Measures of gastrointestinal health and antibiotics use (Figure 4C&D) were obtained through self-administered questionnaires completed by the participants during their initial assessment. For reporting antibiotics use, participants chose from three possible responses (‘not in the past year’, ‘in the past year’, and ‘in the past three months’) which were re-coded into ordinal variables 0, 1, and 2 respectively. Participants chose one of several possible frequencies in response to multiple questions related to gastrointestinal health (abdominal pain, acid reflux, bloating, diarrhea, gas, lack of appetite, nausea) that were re-coded as follows: ‘infrequently/never’=0, ‘once a week or less’=1, ‘more than once a week’=2, ‘daily’=2. The only exception was a question referring to frequency of bowel movement (constipation). In this case, participants chose among the following four answers which were re-coded to the accompanying numerical values: ‘2 or fewer times per week’ = 2, ‘3-6 times per week’=1, ‘1-3 times daily’=0 & ‘4+ times daily’=0. The relationship of each self-reported measure with Shannon diversity was then analyzed independently using linear regression with Shannon and the metabolome-predicted Shannon diversity (mShannon) as the dependent variables and age, sex, and BMI included as covariates. As our cohort consists of self-enrolled participants, response rates of health questionnaires were incomplete; 320 of 399 participants completed the questionnaire on gastrointestinal health and 318 responded to the question regarding antibiotics use. Only responders were used in this particular analysis.

### Analysis of Microbiota and Metabolomic Profiles

A 10-fold cross-validation (CV) implementation of Least Absolute Shrinkage and Selection Operator method (LASSO) was used to predict Shannon diversity from plasma metabolomics data using the Python (version 2.7/3.5+) machine-learning library Scikit-learn. LASSO is a computationally efficient penalized regression method designed to find sparse linear models in large datasets ^32^. All analytes were standardized to mean 0 and unit variance prior to analysis. Ridge regression was also used to compare its performance in predicting Shannon diversity relative to LASSO. Both LASSO and Ridge were fitted with an intercept. A 10-fold CV predict implementation was used to generate a metabolome-predicted Shannon diversity (mShannon) value for each participant. In this approach, each penalized regression model is trained on 90% of the cohort with 10-fold CV, and mShannon diversity is predicted for the 10% of the participants who were not used for model optimization. This process is repeated ten-fold resulting in a ‘test set’ mShannon value for each participant and ten different *β*-coefficients for each metabolite. Using an internal 10-fold CV in each training set and evaluating accuracy of the models using only out-of-sample predictions are meant to control overfitting and provide a conservative estimate of model performance. All metabolites with a non-zero *β*-coefficient in at least one of the ten LASSO models (n=40) were included in further analysis. The R^2^ score was computed by taking the mean of all the R^2^ scores across the 10 out-of-sample predictions. Pearson r was calculated using observed Shannon and mShannon values for the entire cohort.

### Classification Analysis

Eleven metabolites with non-zero β-coefficients across all ten LASSO models generated (Supplementary Table 2) were used to classify participants with low α-diversity using Random forests implemented in Python (version 2.7/3.5+) machine-learning library Scikit-learn. In 3 separate analyses, the cohort was stratified into quartiles based on either Shannon, PD whole tree, or Chao1 diversity metrics. In each analysis, participants in the bottom quartiles were coded as cases (1) while the rest of the cohort was coded as controls (0). Random forests parameters were optimized separately for metabolomics and clinical labs using the GridSearchCV function in Scikit-learn. To confirm the robustness of the identified metabolites as biomarkers for gut diversity, an additional classification was conducted using Shannon diversity as the α-diversity metric in part of the Arivale cohort whose stool samples were analyzed by a different vendor and who were not used for discovery (validation cohort). The same Random forest parameters were used in the validation cohort as the discovery cohort. Mean out-of-sample classification scores and area under the curve (AUC) were calculated using the mean out-of-sample scores across the 10-fold CV.

### Statistical Analysis

Data preprocessing and analysis were both conducted using Python (version 2.7/3.5+). All analytes (proteomics, metabolomics, clinical labs) were scaled and centered prior to analysis. Ordinary Least Square (OLS) linear regression was used to assess the individual relationship between each metabolite retained in at least one of the ten LASSO models (n=40) and Shannon diversity with sex, age, and BMI included as covariates. When assessing the relationships of the strongest metabolite predictors identified using LASSO with Shannon diversity across BMI classes, the cohort was stratified into four groups based on predefined BMI WHO cutoffs. Linear regression was performed on each metabolite individually for each group with Shannon diversity as the dependent variable and sex and age included as covariates. All statistical tests in this study were performed using a two-sided hypothesis. When multiple comparisons were performed, false discovery rate (FDR) was controlled using the method of Benjamini and Hochberg^33^.

## Results

### A small subset of plasma metabolites is strongly predictive of gut microbial α-diversity

In order to investigate the relationship between the host metabolome and gut microbiome diversity, we applied LASSO to baseline plasma metabolomics data of the cohort to generate out-of-sample predictions of Shannon diversity. Of the 659 metabolites measured for each participant in the cohort, only 40 metabolites were retained in the final models (Table 2 & Supplementary Table 2). The small subset of plasma metabolites retained by LASSO was strongly predictive of Shannon diversity, explaining an average 45% of the variance (mean out-of-sample R^2^ score=0.45, Pearson r=0.68 for metabolome-predicted Shannon diversity (mShannon) versus observed Shannon) (Figure 1A). The identified metabolites belonged primarily to the Xenobiotics, Lipid, and Amino Acid superfamilies with a particularly high frequency of phenylalanine/tyrosine metabolites (Figure 1B). While 40 metabolites were retained in at least one of the 10 models generated in the 10-fold CV, only 11 metabolites were retained in all 10 models, and were the most influential in predicting Shannon diversity (Figure 1C, Supplementary Table 2). Ridge regression, which tends to perform better when there is high collinearity among predictor variables and when there is a large number of small effects distributed across the predictor variables ^32^, was outperformed by the LASSO model in our analysis. Ridge with 10-fold CV explained only an average of 35% of variance in Shannon diversity, suggesting that it is indeed a small subset of largely independent plasma metabolites that most strongly reflects gut microbial diversity in our cohort.

**Figure 1:**
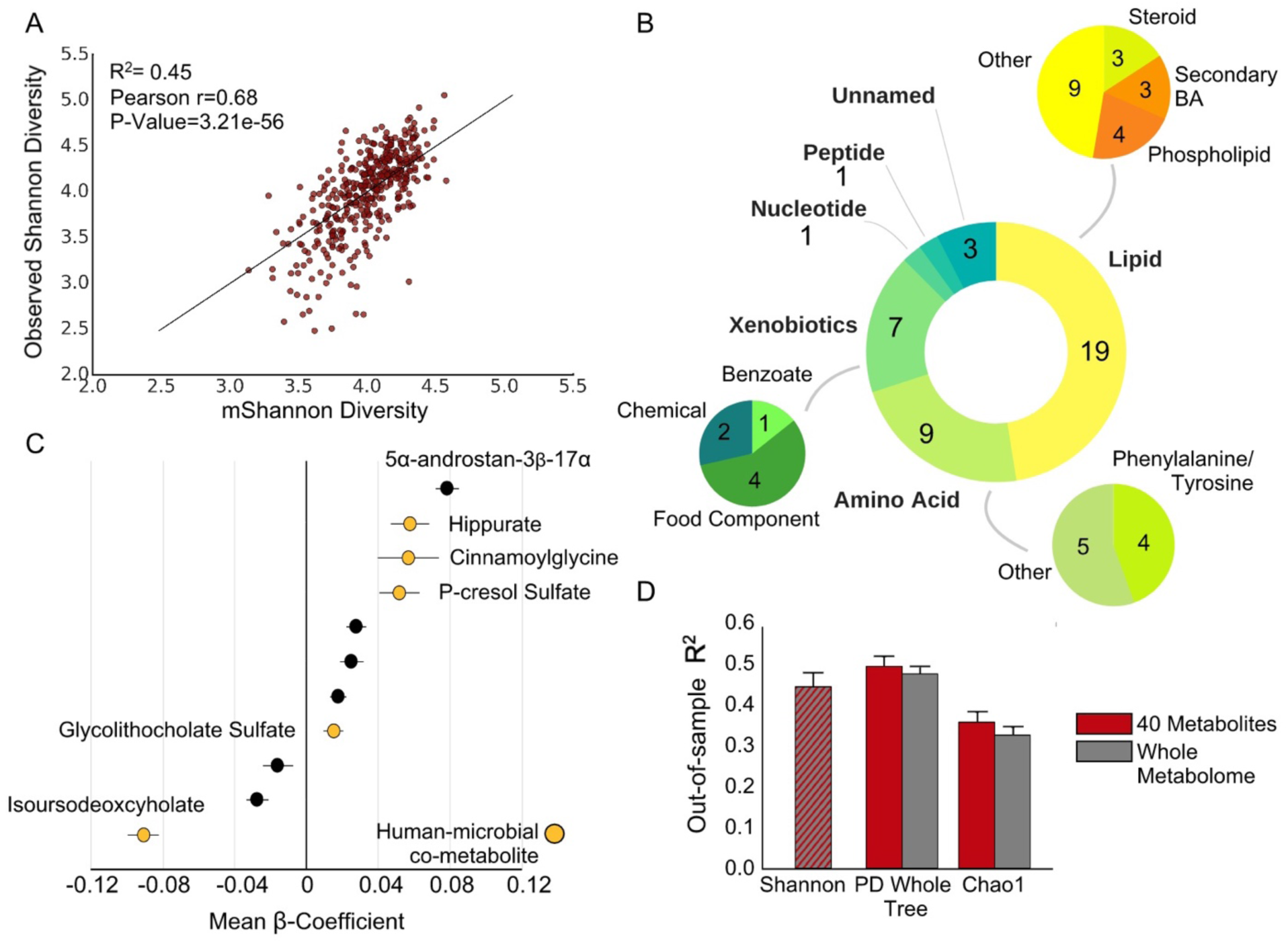
The plasma metabolome is a strong predictor of Shannon diversity. A) A plot of out-of-sample metabolome predicted (mShannon) versus observed Shannon diversity values using LASSO with 10-fold cross-validation. The mean R^2^ across the 10 cross validations, Pearson r of observed versus mShannon values, and corresponding P-Value are shown. B) The super family and subfamily classification of the metabolites (40 total) retained by at least one of the 10 LASSO models used to predict Shannon diversity. Three smaller pie charts correspond to the subfamily classification of metabolites within the Lipid, Amino acid, and Xenobiotics super families. C) A plot of mean β-coefficients corresponding to the 11 metabolites that were retained across all 10 LASSO models. Each coefficient is represented as the mean *β*-coefficient across all 10 models +/- the standard deviation. Yellow dots are labelled and represent *β*-coefficients for human-microbial co-metabolites. Also labelled is the strongest positive predictor 5α-androstan-3β-17α. D) Out-of-sample R^2^ scores of models predicting PD whole tree diversity and Chao1 using either only the 40 metabolites identified (red), or the whole metabolome (grey). Only one bar (dashed red/grey) is shown for Shannon diversity, since the 40 metabolites were identified using LASSO on the whole metabolome. Values are presented as mean out-of-sample R^2^ score (n=10) +/- standard error of the mean. *Abbreviations: 5α-androstan-3β-17α: 5α-androstan-3β-17α-diol disulfate; BA: bile acids*.

**Table 2:**
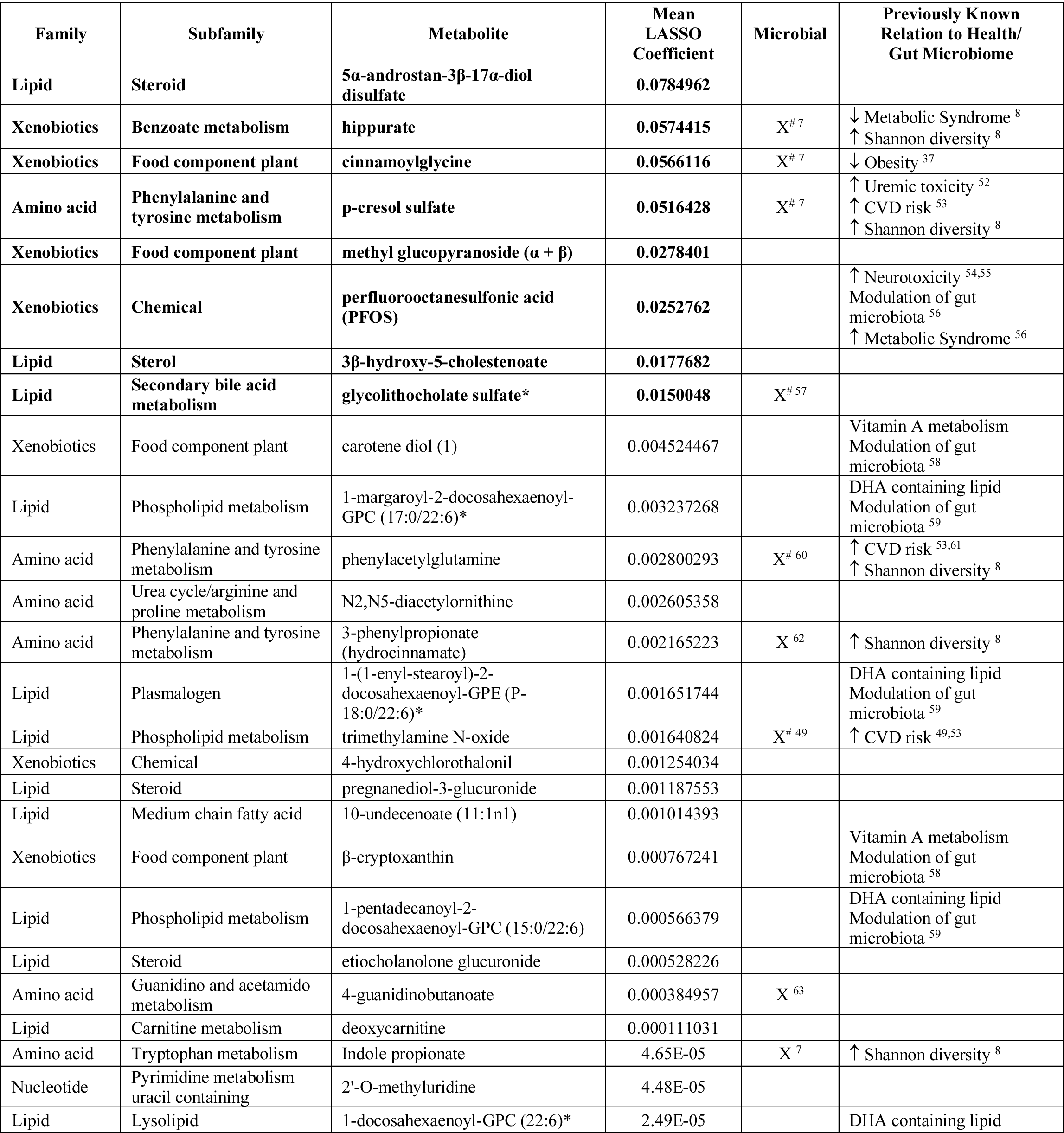
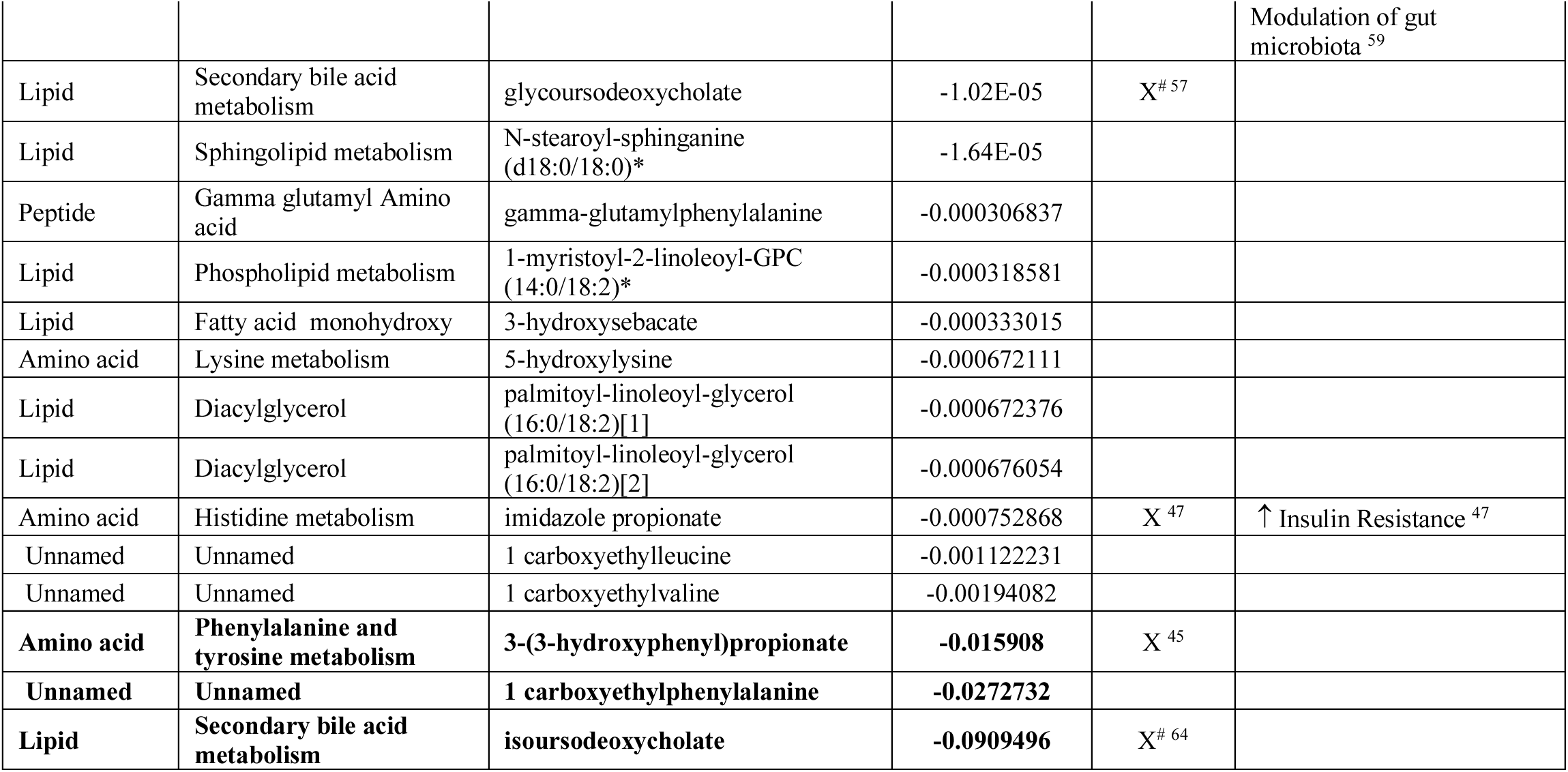
Relationships of the 40 identified metabolites with human health and gut microbiome structure. The family, subfamily, as well as mean *β*-coefficient across the 10 LASSO models generated are provided. ^‘#’^ indicates a human microbial co-metabolite. The 11 metabolites that were retained across all 10 LASSO models are bolded.

We investigated potential collinearity among the 40 metabolites identified through LASSO. Of the 1560 metabolite-metabolite comparisons, only six were identified to be highly collinear (|r|>0.80) (Supplementary Figure 1 A&B). These included correlations among three 1-carboxyethyl amino acids (leucine, phenylalanine, valine). Of the three, 1-carboxyethylphenylalanine was retained across all 10 models while the other two carboxyethyl compounds were retained in two (valine) and three (leucine) models. The low collinearity among predictor variables is consistent with the way LASSO generates a regression model. When analytes are highly collinear, LASSO will retain one of the analytes in the model while pushing the *β*-coefficients of the other analytes toward zero ^32^.

We further assessed the capacity of the 40 identified metabolites to predict other metrics of gut microbial α-diversity including PD whole tree and Chao1 (Figure 1D). The first of these two metrics incorporates phylogenetic differences in the diversity score, while the second is a measure of species richness. A LASSO model fitted using 10-fold CV with only the 40 identified metabolites explained an average 50% of out-of-sample variance in PD whole tree diversity and 36% in Chao1. Importantly, using the whole plasma metabolome did not improve the predictive capacity for either of the two metrics, confirming that the small subset of metabolites identified is sufficient to capture majority of the explainable variability in gut microbial α-diversity.

Many of the strongest identified predictors of Shannon diversity in the models generated using LASSO were human-microbial co-metabolites: metabolites either initially synthesized by the host and then subsequently metabolized by the microbiome (e.g. bile acids) or vice versa (e.g. hippurate) (Figure 1C and Table 2). Our results confirm previously identified metabolites such as hippurate, and p-cresol sulfate ^8^, as well as identify novel candidate biomarkers of Shannon diversity (Figure 1C, Table 2). In particular, the testosterone metabolite 5α-androstan-3*β*-17α-diol disulfate (5α-androstan-3*β*-17α) had the highest mean positive β-coefficient across all LASSO models generated (Figure 1C). Plasma concentrations of 5α-androstan-3*β*-17α were significantly higher in men than in women, while no significant sex dependent differences were observed in Shannon diversity (linear models adjusted for age and BMI) (Figure 2A&B). There was a significant positive association between Shannon diversity and 5α-androstan-3*β*-17α for both men and women. While there appeared to be a variable relationship between Shannon diversity and 5α-androstan-3*β*-17α (Figure 2C) across sex, a sex by 5α-androstan-3*β*-17α interaction term in a regression model adjusted for age and BMI did not reach significance (*β*(*men*5α-androstan-3β-17α*)(95%CI) =0.072(−0.034, 0.18), P-Value=0.18).

**Figure 2:**
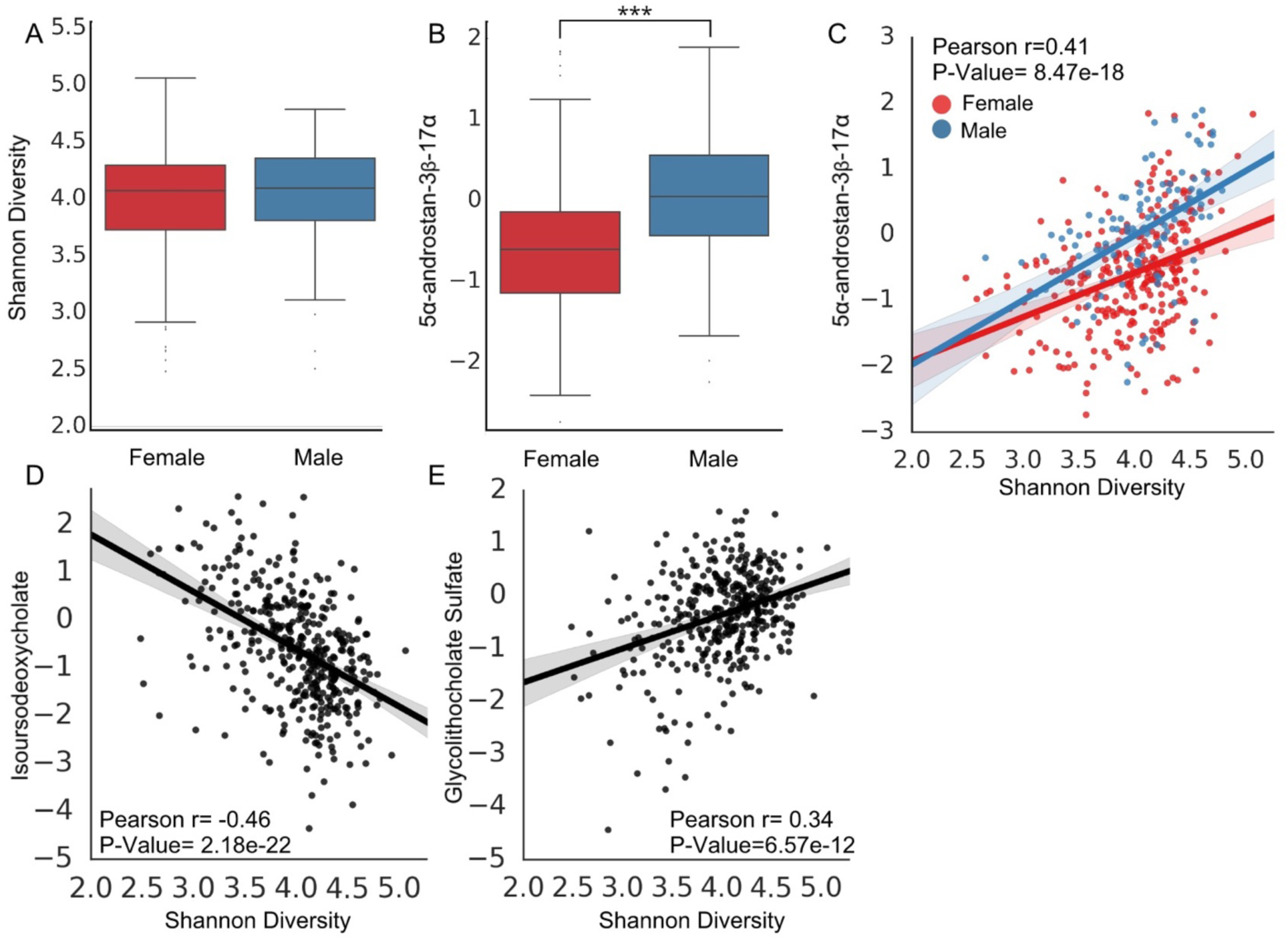
5α-androstan-3*β*-17α correlates with Shannon diversity in males and females. A) Shannon diversity is not significantly different across sex. B) 5α-androstan-3*β*-17α blood concentration is higher in men (n=111) than women (n=288), adjusted for age and BMI. (P-Value=3.79e-13). C) 5α-androstan-3*β*-17α is positively associated with Shannon diversity in both males and females. D&E) Secondary bile acids retained by LASSO in the prediction model show opposite association with Shannon diversity. *Abbreviations: 5α-androstan-3β-17α: 5α-androstan-3β-17α-diol disulfate*.

Two secondary bile acids, isoursodeoxycholate and glycolithocholate sulfate, were also retained in all 10 LASSO models (Supplementary Table 2). Interestingly, they demonstrated opposite associations with Shannon diversity in our cohort (Figure 2D&E). When the top 11 metabolites from LASSO were correlated with bacterial genera, the two bile acids consistently demonstrated opposite associations; glycolithocholate sulfate was anticorrelated with the most abundant Bacteroidetes genus, *Bacteroides*, while isoursodeoxycholate demonstrated a positive association. The opposite was true for several Firmicutes genera (Figure 3 & Supplementary Figure 2). *Bacteroides* was also anti-correlated with the strongest positive predictor 5α-androstan-3*β*-17α, indicating that increasing dominance of this taxon is related to reduced Shannon diversity in our cohort.

**Figure 3:**
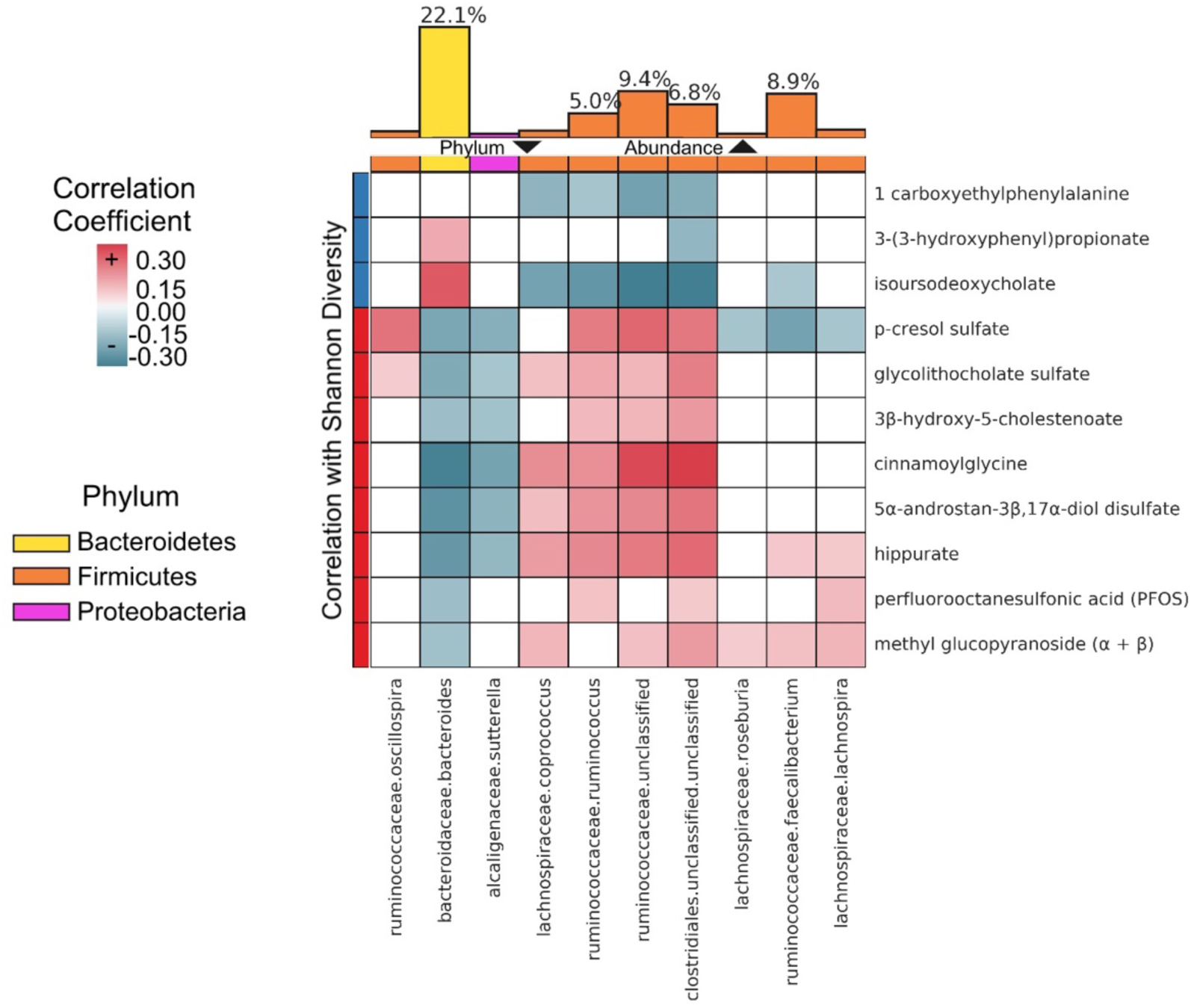
Significant Spearman correlations of each of the 11 metabolites retained by all 10 LASSO models (rows) with the 10 most abundant microbiome genera (columns). Top color row labels the phylum for each genus. Left color column labels the sign of the correlation between the metabolite and Shannon diversity (blue - negative correlation, red - positive correlation). The top bar graph represents the median fractional abundance of each genus across the cohort, with bars colored by phylum. Non-significant correlations are colored in white (FDR<0.05).

The relationship between each metabolite that had a non-zero *β*-coefficient in at least one of the 10 LASSO models generated and Shannon diversity was next independently assessed using linear regression with sex, age, and BMI included as covariates. Of the 40 identified metabolites, 35 were significantly associated with Shannon diversity (FDR<0.05) (Supplementary Table 1). Shannon diversity was also regressed independently against each metabolite of the 40 identified using LASSO. Cinnamoylglycine alone explained the greatest percent of variance (25.6%) of all metabolites (Figure 4A). An additional 8 metabolites explained over 10% of variance in Shannon diversity each. However, no single metabolite explained a similar percent of variance as LASSO predicted mShannon, demonstrating that a combination of plasma metabolites is substantially more reflective of Shannon diversity than any one metabolite alone.

**Figure 4:**
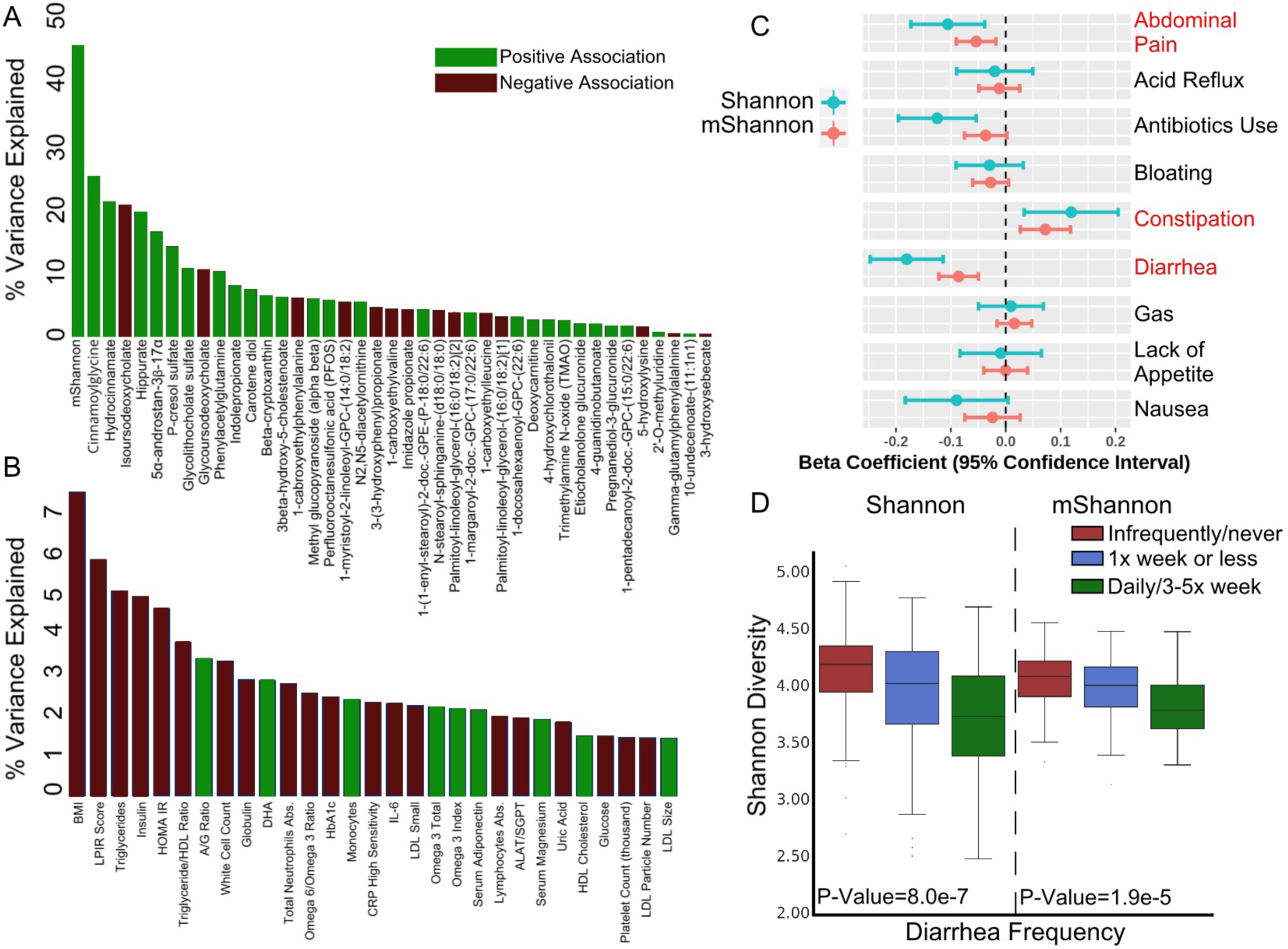
Reflection of Shannon diversity in clinical laboratory tests and the host metabolome. A) The percent of variance in Shannon diversity explained by each plasma metabolite retained by at least one of the 10 LASSO models generated individually. mShannon is included for comparison. B) The percent of variance in Shannon diversity explained by each blood clinical analyte significantly associated with Shannon (FDR<0.05) individually. BMI is included for comparison. Red bars correspond to a negative, while green bars correspond to a positive, β-coefficient for each analyte. C) β-coefficients and 95% confidence intervals for each metric of gastrointestinal health in a model with Shannon or mShannon Diversity as the dependent variable and sex, age, and BMI included as covariates. Each metric of gastrointestinal health was coded on a three-point scale (see Methods). Significant associations for both Shannon and mShannon are highlighted in red. D) Box plots for Shannon and mShannon diversity stratified across self-reported frequency of diarrhea (Infrequently/never n=150, 1x week or less n=127, Daily/3-5x a week n=43).

As mentioned previously, only 11 of the 40 metabolites identified were retained in all 10 LASSO models (Figure 1C). These metabolites were next used to classify participants with low α-diversity (bottom quartile) using a 10-fold CV implementation of Random forests. The 11 metabolites alone were able to classify participants based on not only Shannon (mean Sensitivity=0.72, Specificity=0.90, Precision=0.72) but also other metrics of α-diversity (PD whole tree: mean Sensitivity=0.68, Specificity=0.89, Precision=0.67; Chao1: mean Sensitivity=0.64, Specificity=0.88, Precision=0.65). Together these results indicate a potential use for the 11 identified metabolites as biomarkers for gut microbial diversity.

### Clinical laboratory tests are not predictive of gut microbial diversity

To date, blood clinical laboratory tests are still the most commonly used molecular metric for routinely evaluating the health state of an individual. However, it is unclear whether or how effectively laboratory tests may capture the structure of the gut microbiome. To assess whether common clinical labs reflect the state of the gut microbiome, we utilized the same methods as with metabolomics in order to predict Shannon diversity from a wide panel of clinical analytes. A set of 77 clinical labs could not accurately predict individual Shannon diversity scores (mean LASSO R^2^= 0.01, mean Ridge R^2^=0.05). The weak association of clinical labs and Shannon diversity was further confirmed when linear regression was used to assess the relationship of each clinical lab and Shannon diversity independently. Of 77 clinical labs, 28 were significantly associated with Shannon diversity (FDR<0.05) (Figure 4B, Supplementary Table 3). However, no clinical analyte explained more than 6% of variance in Shannon diversity alone, with markers of metabolic health (lipoprotein insulin resistance (LPIR) Score, triglycerides, insulin) ranking the highest of all analytes (Figure 4B). By comparison, BMI alone explained 7.5% of variance in Shannon diversity, explaining a greater percent of variance in Shannon diversity than any single clinical analyte. When individual clinical analyte models were adjusted for sex, age and BMI, only LPIR Score and blood triglycerides remained significantly associated with Shannon diversity (Supplementary Table 3) (FDR<0.05). Concordantly, a Random forests classification model based on clinical labs optimized to identify participants with low Shannon diversity (bottom quartile) demonstrated low performance (sensitivity=0.50, specificity=0.80, precision=0.45) (Supplementary Figure 3).

### Metabolic health related blood proteins are associated with Shannon diversity

In addition to clinical labs and metabolomics, 263 unique blood proteins were measured for 262 of 399 participants in our cohort. Using linear regression and adjusting for multiple hypothesis testing, 41 of the 263 proteins were identified to be significantly associated with Shannon diversity (FDR<0.05) (Supplementary Table 4). Similar to results of the clinical labs (Figure 4B), the most strongly associated analytes were proteins involved in metabolic health including Paraoxonase 3 (PON3) (*β*(95%CI)=0.13 (0.084, 0.18), P-Value=4.7e-08), Leptin (LEP) (*β*(95%CI)= −0.14 (−0.20,-0.088), P-Value=5.1e-07) and Low-Density Lipoprotein Receptor (LDLR) (*β*(95%CI)= −0.12 (−0.17,-0.064), P-Value=2.9e-05). PON3 is an enzyme that associates with HDL and may prevent oxidation of LDL ^34^, while LEP is an adipocyte derived hormone that is elevated in obesity ^35^. After adjusting for covariates, only three proteins remained significantly associated with gut microbial diversity, including LDLR, hydroxyacid oxidase 1 (HAO1), and serine protease 8 (PRSS8) (Supplementary Table 4). Finally, the same penalized regression methods were used as with metabolomics in order to evaluate how strongly blood proteins measured reflect Shannon diversity. The plasma proteome was more predictive of Shannon than clinical labs, explaining an average 13% of variance (mean LASSO R^2^ Score= 0.11, mean Ridge R^2^ Score=0.13). However, the prediction was still substantially weaker than that of the plasma metabolome.

### mShannon reflects perturbations associated with gastrointestinal health

Given that the plasma metabolome was the strongest predictor of gut microbiome structure, we moved forward with our mShannon predictions to assess whether host physiology reflects perturbations in the gut microbiota associated with several lifestyle and medical factors reported previously. Using information from lifestyle and health questionnaires completed by Arivale participants, Shannon diversity was regressed against self-reported measures of gastrointestinal health as well as antibiotics use, adjusting for covariates (see Methods). Several factors significantly associated with major perturbations in Shannon diversity were also reflected in the host’s metabolome, as shown by significant β-coefficients in models predicting both Shannon and mShannon (Figure 4C). In particular, reported frequency of diarrhea and abdominal pain demonstrated a strong negative association with Shannon diversity, which was reflected in mShannon as well (Figure 4C&D). Antibiotics use was significantly associated with lower Shannon diversity, with the same trend observed in mShannon (P-value=0.06). Lower frequency of bowel movements (constipation) was positively associated with both Shannon diversity and mShannon (Figure 4C). Together these results suggest that events associated with major changes in gut microbial structure and gut health are strongly reflected in the host metabolome.

### Plasma metabolites associated with Shannon diversity vary across BMI classes

Obesity has been consistently linked to metabolic perturbations ^36^ as well substantial changes in the blood metabolome ^37^. Additionally, some but not all studies have reported a decrease in Shannon diversity in obese versus normal weight individuals ^19^. To explore the relationship between Shannon diversity, obesity and the plasma metabolome further, the cohort was stratified based on BMI into four groups (Table 1). Because the risks associated with obesity vary by the degree of BMI increase, obesity is typically divided into three classes (Class I = 30-34.9 kg/m^2^, Class II = 35-39.9 kg/m^2^, and Class III ≥40 kg/m^2^). For this analysis we combined Class II and class III obesity but looked at Class I (“low-risk obesity”) separately. ^38,39^ All three metrics of gut α-diversity were significantly decreased with increasing BMI, with participants in the obese II/III class demonstrating the lowest diversity scores of all groups (Table 1). In particular, for both Chao1 and PD whole tree diversity, participants in the obese II/III group had a significantly lower score than all other BMI classes, including obese I. Molecular markers of health were progressively worse across BMI classes, however markers of inflammation, such as Interleukin-6 and C-Reactive Protein, were particularly elevated in the obese II/III group (Table 1, Figure 5A).

**Figure 5:**
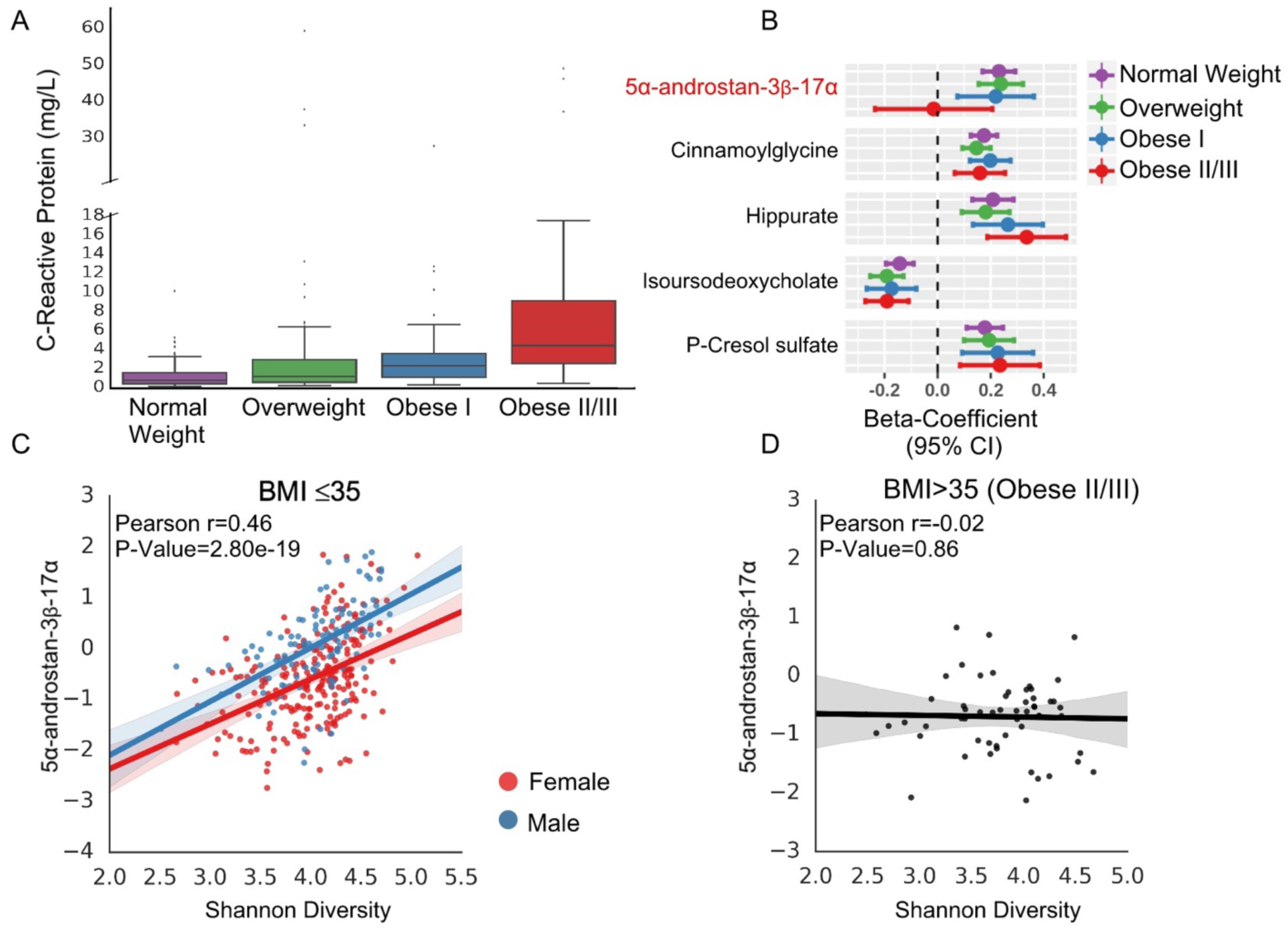
The gut microbiome-host metabolome relationship is perturbed under severe obesity. A) CRP levels across BMI classes in the cohort. B) β-coefficients for each of the strongest 5 metabolite predictors of Shannon diversity identified through LASSO. The cohort was stratified based on BMI class and Shannon was regressed against each metabolite individually with sex and age included as covariates. C) Scatter plot representing the relationship between 5α-androstan-3*β*-17α and Shannon diversity in participants with BMI ≤ 35. Regression line was fitted for males and females, separately. D) Scatter plot representing the relationship between 5α-androstan-3*β*-17α and Shannon diversity in obese II/III participants. A single regression line was fitted due to the small number of men in the group (n=4). Pearson r and corresponding P-value are shown. *Abbreviations: 5α-androstan-3β-17α: 5α-androstan-3β-17α-diol disulfate*.

To evaluate whether there is variability in the reflection of Shannon diversity in the plasma metabolome across BMI classes, the relationship between Shannon and the five strongest metabolite predictors retained by all 10 LASSO models (greatest absolute value of mean *β*-coefficients) was assessed within each BMI class, adjusted for age and sex (Figure 5B). While the *β*-coefficients were similar across all BMI classes for most metabolites, 5α-androstan-3*β*-17α demonstrated no significant association with Shannon diversity in the obese II/III group, yet a significant positive association among all other participants (Figure 5C&D). When the analysis was extended to all 11 metabolites that were retained by the 10 LASSO models generated (Supplementary Figure 4A), other differences emerged, including a significant relationship between Shannon and the bile acid intermediate 3*β*-hydroxy-5-cholestenoate in the obese I class but not all other BMI classes, and a stronger relationship between Shannon and perfluorooctanesulfonic acid (PFOS) in the obese II/III class relative to all other groups. PFOS alone explained 22.4% (Pearson r=0.48, P-Value=2.0e-04) of variance in Shannon in obese II/III participants, while no significant relationship was detected in normal weight participants independently (Pearson r=0.09, P-Value=0.30) and after adjusting for covariates (*β*(95%CI)=0.04 (−0.036,0.11), P-Value=0.32) (Supplementary Figure 3B). Only a weak association between PFOS and Shannon was observed when all non-obese II/III participants were pooled together (Pearson r=0.20, P-Value=2.2e-04), highlighting the variable reflection of gut α-diversity in the host metabolome across the BMI spectrum.

### The host metabolome/gut microbiome relationship is consistent in a validation cohort

To confirm our findings, major observations from the discovery cohort were explored in a separate group of Arivale participants who joined the program at an earlier date and whose blood and microbiome samples were analyzed by different vendors (clinical labs: LCA (38%), Quest Diagnostics (62%), microbiome= Second Genome (100%), N=540). Of the 35 metabolites significantly associated with Shannon diversity after adjusting for covariates and multiple hypothesis testing in the discovery cohort, 29 were confirmed in the validation cohort (Supplementary Table 2). Similarly, 66% of the significant microbiome genera-plasma metabolite correlations identified in our discovery cohort were confirmed in the validation set (FDR<0.05) (Supplementary Figure 5&6). A LASSO model using only the 40 metabolites identified in the discovery cohort was further fitted to predict Shannon diversity in the validation cohort using 10-fold CV. Out-of-sample prediction accuracy for the 40-metabolite LASSO model was lower but similar to that in the discovery cohort (mean R^2^=0.34, Pearson r=0.60, Figure 6A&C). Additionally, *β*-coefficients for the 11 metabolites that were retained across all LASSO models in the discovery cohort were compared to the mean *β*-coefficients fitted on the validation cohort using 10-fold CV, showing a strong correlation between models (Pearson r=0.94, P-value= 2.05e-05, Figure 6B). The 40 metabolites alone explained majority of the variance in Shannon diversity captured by the blood metabolome in the validation set. Generating a separate LASSO model using the whole metabolome (659 metabolites) improved the predictive capacity of the model only marginally (mean R^2^=0.38, Figure 6C), with the mean difference in performance across the 10-fold CV not being significantly different between the 40 metabolite and the whole metabolome model (P-value=0.30). The whole metabolome LASSO model in the validation cohort retained 58 metabolites across the 10 CVs. Of the 58, 15 overlapped with the discovery cohort set of 40 metabolites (Supplementary Table 2). This is unsurprising given the various demographic differences across the two cohorts (Supplementary Table 2) and the way in which LASSO retains analytes in the model. Despite the moderate overlap between metabolites retained independently by applying LASSO in the discovery and validation sets, mean *β*-coefficients across the 10-CVs for the 11 strongest metabolite predictors still showed strong correlation between models (spearman ρ=0.91, P-value=9.2e-05, Pearson r=0.90, P-value=1.9e-4). The predictive capacities of clinical labs and proteomics were also investigated in the validation cohort. Consistent with our initial findings, clinical labs demonstrated little to no correspondence with Shannon diversity (Figure 6C). Only 176 of the 540 participants in the validation set had available proteomics data. In this sub-group of the validation cohort, we were unable to confirm the moderate capacity of the blood proteome to predict Shannon diversity, with the mean variance explained lower than clinical labs (Figure 6C). Finally, classification of participants with low Shannon diversity (bottom quartile) using the 11 strongest Shannon diversity predictors identified in the discovery cohort was investigated in the validation set where similar Sensitivity (0.65), Specificity (0.87), and Precision (0.64) were achieved (Figure 6D&E). Collectively these findings increase our confidence in the strong relationship between the 40, and in particular the core 11, identified metabolites and gut microbiome diversity.

**Figure 6:**
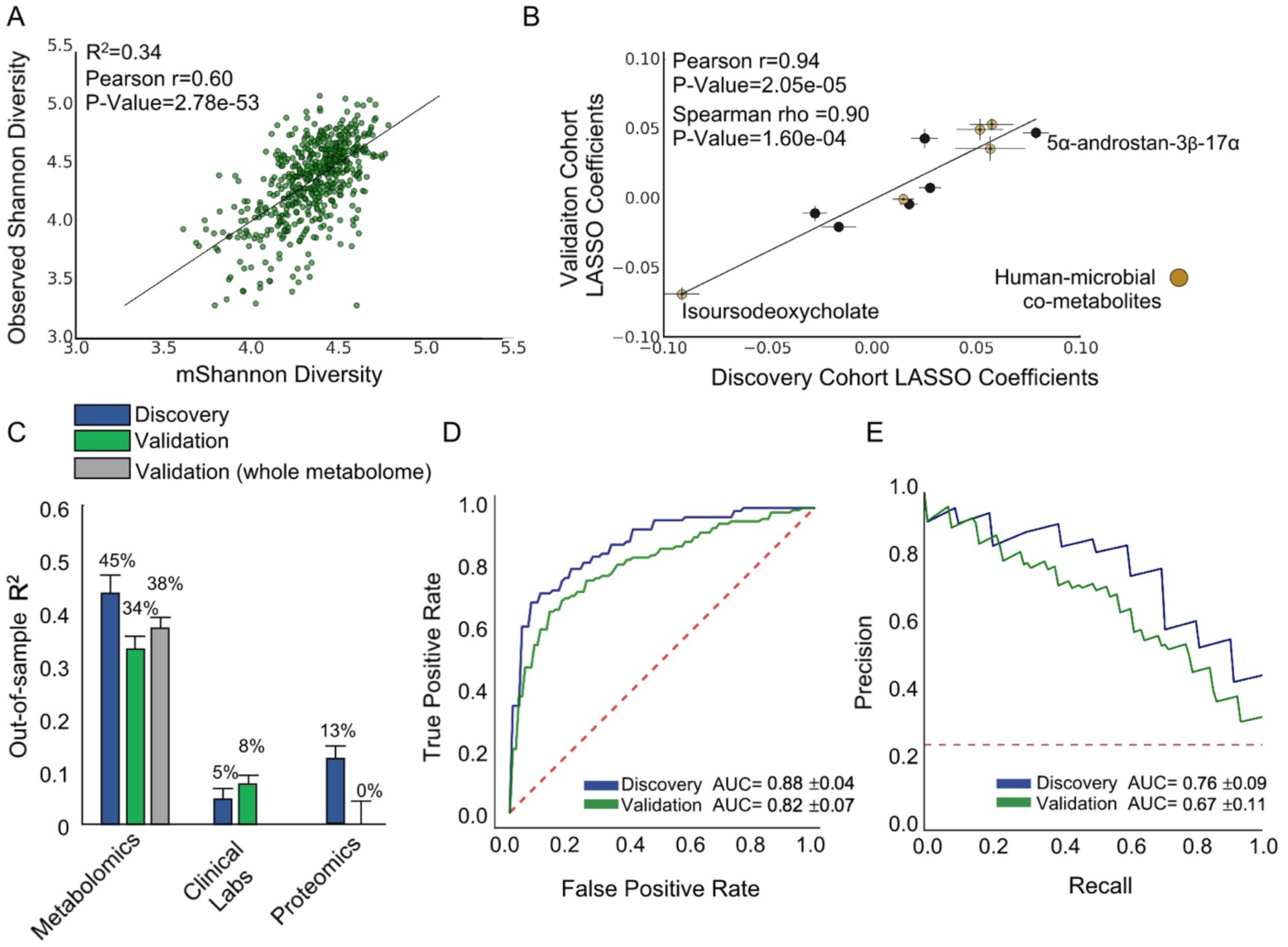
Gut-microbiome/host metabolome relationship is consistent in a validation cohort. A) Scatter plot of observed Shannon diversity and mShannon diversity predicted in the validation cohort using LASSO with only the 40 identified metabolites from the discovery cohort. Mean out-of-sample R^2^ across the 10 cross validations and Pearson r for mShannon versus observed Shannon are shown. B) Mean LASSO *β*-coefficients for the 11 strongest metabolite predictors of mShannon diversity from the discovery cohort strongly correlate with the Mean LASSO *β*-coefficients for the same metabolites optimized in the validation cohort. C) Percent of variance explained using penalized regression across omics platforms in the discovery and validation cohorts. Values are presented as mean R^2^ across 10-fold CV +/-standard error of the mean. The grey bar for metabolomics corresponds to a model fitted using all 659 metabolites, while the green bar corresponds to a model fitted using only the 40 metabolites identified in the discovery cohort. D) Receiver Operator Characteristic curves classifying participants with low Shannon diversity (bottom quartile) for the discovery and validation cohorts. E) Precision-Recall curves classifying participants with low Shannon Diversity (bottom quartile) for the discovery and validation cohorts. Mean area under the curve (AUC) values across 10-fold CV +/-standard deviation are shown.

We next explored key differences in the host metabolome/gut microbiome relationship across obesity identified in the discovery cohort. However, the number of participants in the obese II/III group in the validation set was substantially lower than in the discovery cohort (n=34), limiting our statistical power. Consistent with our initial findings, PFOS was weakly positively associated with Shannon diversity in normal weight participants (BMI<25) (Pearson r=0.17, P-Value=0.01). This relationship was once again stronger in obese II/III participants (Pearson r=0.35, P-Value=0.04), though to a lesser extent than in the discovery cohort (Supplementary Figure 4C). After adjusting for covariates, PFOS was no longer significantly associated with Shannon in the obese II/III group (*β*(95%CI)= 0.11 (−0.04,0.24), P-Value=0.15). The sex steroid hormone 5α-androstan-3β-17α demonstrated a significant positive correlation with Shannon diversity across participants with BMI ≤35, consistent with the discovery cohort (Supplementary Figure 4C). The metabolite also remained one of the strongest positive predictors in the LASSO models generated (Figure 6B). However, no significant relationship was observed in obese II/III participants between Shannon diversity and 5α-androstan-3β-17α independently (Pearson r=−0.01, P-Value= 0.95) and after adjusting for covariates (*β*(95%CI)=0.14 (−0.02,0.30), P-Value=0.08) (Supplementary Figure 3D). To increase statistical power, obese II/III participants were pooled together from both the discovery and validation cohorts (n=90) and the relationship between 5α-androstan-3β-17α and Shannon diversity was assessed using linear regression, adjusting for sex, age, and microbiome vendor. Even with increased power, there was no significant association between 5α-androstan-3β-17α and Shannon (*β*(95%CI)=0.03 (−0.10,0.15), P-Value=0.70), providing further evidence for a lack of relationship of this metabolite with Shannon diversity under severe obesity.

## Discussion

The goal of this study was to examine the relationship between gut microbiome α-diversity and a broad set of analytes measured in the blood. We generated and analyzed baseline data from a cohort of hundreds of individuals participating in a wellness program, including numerous blood measurements (metabolomes, proteomes, and clinical lab tests) and fecal microbiomes taken within 21 days of each other. The key findings of this study were: (1) 40 plasma metabolites, but not a wide clinical lab panel or a set of 263 proteins, were able to predict gut microbiome Shannon diversity in a cohort of predominantly healthy US adults; (2) specific metabolites associated with Shannon diversity in our study are also related to human health, such as the microbial metabolites trimethylamine N-oxide (TMAO), imidazole propionate and p-cresol sulfate; (3) Shannon diversity predictions accurately reflected changes in gut microbial diversity associated with gastrointestinal health, demonstrating a strong connection between gut microbiome structure and host physiology; and (4) there was a variable relationship between specific metabolites and Shannon diversity across the BMI spectrum, with particular disruption to this relationship in severe (class II/III) obesity. Collectively, our results provide novel insight into the intimate relationship between host physiology and the gut microbiome and characterize the host metabolome as an important interface between the gut ecosystem and human health.

The weak capacity of clinical labs to predict Shannon diversity scores and accurately classify participants with low gut microbial diversity highlights the need for more refined molecular biomarkers for assessing gut microbiome health. The strong imprint of Shannon diversity in the plasma metabolome is perhaps unsurprising, given that our untargeted metabolomics panel captures many metabolites of microbial origin. The advantage of analyzing the plasma metabolome as a marker of gut microbiome structure is the ease of sample collection and processing, as well as the biological relevance of metabolomics obtained from the blood. Unlike fecal metabolites that have been previously shown to be strongly related to gut microbial composition ^6^, the plasma metabolome captures analytes that have greater potential to exert a biological effect in the host. We anticipate that many of the metabolites identified in our analysis can be investigated further to better understand the impact of the gut microbiome on human health.

A more diverse microbiome is often assumed to be a healthier microbiome ^40–42^. Consistent with this assumption, several metabolites with positive *β*-coefficients in the LASSO models generated to predict Shannon diversity in our study have been previously linked to microbial metabolism of health-promoting polyphenolic compounds (e.g. benzoate that is conjugated in the liver to hippurate, as well as hydrocinnamate) ^43–45^. Polyphenols are a diverse group of phytochemical compounds found in fruits, vegetables, and cereals as well as coffee, tea and red wine ^46^. The positive association between polyphenolic microbial metabolites and Shannon diversity suggests a diverse microbiome may reflect a polyphenol rich diet. Similarly, the potentially detrimental microbial metabolite imidazole propionate was negatively associated with Shannon diversity across both the discovery and validation cohorts. A recent study demonstrated that imidazole propionate is synthesized in greater abundance in the gut microbiota of diabetic patients, and that it is able to impair insulin signaling in animal models ^47^. Its negative association with α-diversity in our study supports the notion that host health increases with increased microbial diversity. However, our results also suggest that there may be an optimal range of α-diversity, rather than a monotonic relationship between microbiome diversity and human health. For example, p-cresol sulfate was positively correlated with Shannon diversity and is a potentially toxic uremic compound formed from the putrefaction of undigested dietary proteins by colonic bacteria and subsequent modification by the liver ^48^. Similarly, the human-microbial co-metabolite TMAO was positively associated with Shannon diversity in both the discovery and validation cohorts, independent of age, sex, and BMI. TMAO is associated with diets high in red meat. The negative effects of TMAO on health are still emerging, but growing evidence suggests it may promote cardiovascular disease ^26,49^. Fermentation of dietary protein accelerates when dietary fiber is exhausted in the stool. Thus, higher levels of protein fermentation are found in people on high-protein/low-fiber diets and in people suffering from constipation ^50^. Consistently, participants who reported constipation in our study also showed higher metabolome-predicted Shannon diversity. We suggest that, above a threshold, higher α-diversity may be associated with unhealthy blood levels of particular microbial metabolites. Alternatively, we found that very low Shannon and mShannon diversity scores were associated with participant-reported diarrhea and abdominal pain. Together, these results provide evidence for the existence of a ‘Goldilocks Zone’ for gut microbiome diversity.

The relationship between the human gut microbiome and obesity has been inconsistent in the literature ^19^. Previous work has often grouped participants into just two categories: healthy and obese. In this study, we stratified participants into multiple BMI classes. We hypothesized that participants who are in the obese II or III class, which are both associated with a higher risk of a spectrum of diseases compared to obese I individuals ^38,39^, will show a greater disruption to their gut microbiome. Through this refined stratification, we were able to demonstrate that the associations between the plasma metabolome and microbiome diversity were qualitatively different across different BMI classes. In particular, the sex steroid 5α-androstan-3*β*-17α emerged as one of the strongest positive predictors of Shannon diversity in both the discovery and validation cohorts, while demonstrating no significant correlation with Shannon diversity in obese II/III individuals. Previous studies have shown that the gut microbiome is capable of altering sex steroid hormone levels (testosterone) in the host ^4^, providing a potential direct link between the identified testosterone metabolite and the gut microbiota. The lack of relationship between gut diversity and 5α-androstan-3*β*-17α in obese II/III participants may have potential health implications. Alternatively, the known impact of severe obesity on sex steroids may be playing a role in the differential findings in obese II/III participants ^51^. However, further studies are needed to fully elucidate the relationship between the identified hormone, the gut microbiome, and metabolic health. The environmental pollutant PFOS showed a positive association with Shannon diversity in obese II/III participants, but no significant association in normal weight individuals in the discovery cohort and a weak association in the validation cohort. These differences highlight the importance of refining our definitions of obesity when investigating the impact of the gut microbiome on health and also stress context when assessing the relationship between microbial diversity and host physiology.

Despite the hundreds of associations between the gut microbiome and disease that have been identified over the last decade, we still do not yet understand what constitutes a ‘healthy’ microbiome-host relationship. It is likely that the connection between microbiome community structure and human health is highly contextual and complex, depending on diet, behavior, exposure to pathogens, history of antibiotic exposure, genetics, and other myriad factors. Our work integrates multi-omics data from a large cohort of individual people to provide a path forward for elucidating the properties of a ‘healthy’ microbiome-host interaction. We provide new insight based on the metabolic states of individual hosts and identify severe obesity as a crucial factor for stratifying patient populations. While we cannot establish causality, this work advances our knowledge of how host metabolism and microbial community structure are inter-connected. Understanding this interplay in a multi-omics context will enable future experimental work and the development of personalized, multimodal interventions aimed at promoting wellness and treating disease.

### Institutional Review Board Approval for the Study

Procedures for the current study were run under the Western Institutional Review Board (WIRB) with Institutional Review Board (IRB) Study Number 20170658 at the Institute for Systems Biology and 1178906 at Arivale (both in Seattle, WA).

### Data and Code Availability

The model summary statistics for all metabolites, proteins and clinical labs analyzed are available to download in Supplementary Tables 2-4. Researchers can request to access the full de-identified dataset of microbiome and metabolomics by signing the Arivale data access agreement: arivale.com/legal/data-access. The necessary packages and code will be made available at https://github.com/PriceLab/ShannonMets.

## ACKNOWLEDGMENTS

We thank Christian Diener, Anat Tzimmer and Max Robinson for helpful discussions.

## AUTHOR CONTRIBUTIONS

T.W., N.R., S.M.G., L.H. and N.D.P. conceived of the study. T.W., N.R., J.C.E, G.S.O, S.M.G., L.H., and N.D.P., participated in study design. T.W, N.R, A.M., O.T.M. processed the data and performed computational analysis. A.T.M, O.T.M., and J.L. managed logistics of data collection and integration. T.W., N.R., S.M.G and N.D.P were the primary writers of the paper, with contributions from all authors. All authors read and approved the final manuscript.

## FUNDING

This research was supported by The M.J. Murdock Charitable Trust (L.H., N.D.P.), Arivale, and a generous gift from Carole Ellison. S.M.G. was supported by a Washington Research Foundation Distinguished Investigator Award.

## Competing financial interests

L.H. and N.D.P. are co-founders of Arivale (where these data come from) and hold stock in the company. N.D.P. is on the Arivale board of directors; L.H. is chair of and G.S.O. is a member of Arivale’s scientific advisory board. A.T.M., O.M. and J.L. are employees of Arivale and have stock options in the company, as do G.S.O. and J.C.E.

## Supplementary Tables

**Supplementary Table 1.** Validation cohort characteristics overall and by BMI class. Values with different superscript letters are significantly different (P<0.05). ** indicates P<0.001 and *** P<0.0001. NS – not significantly different between cohorts.

|  | **Normal**<br>**Weight** | **Overweight** | **Obese I** | **Obese II/III** | **Total** | **Change**<br>**relative to**<br>**discovery**<br>**cohort** |
| --- | --- | --- | --- | --- | --- | --- |
|  | **(n=256)** | **(n=167)** | **(n=83)** | **(n=34)** | **(N=540)** |  |
| **Mean Age** | 48.4 | 50.9 | 52.7 | 49.3 | 49.9 | +2.9 yrs** |
| **(SD)** | (11.9)<sup>a</sup> | (13.1)<sup>b</sup> | (11.6)<sup>b</sup> | (10.9)<sup>a,b</sup> | (12.3) |  |
| **Male (%)** | 39.8<sup>a</sup> | 59.9<sup>b</sup> | 48.2<sup>a,b</sup> | 38.2<sup>a</sup> | 47.2 | +19.6%*** |
| **Non-white (%)** | 23.4 | 18.6 | 24.1 | 23.5 | 22 | NS |
| **Median BMI** | 22.8<sup>a</sup> | 27.0<sup>b</sup> | 31.4<sup>c</sup> | 37.1<sup>d</sup> | 25.2 | -1.6 *** |
| **(IQR)** | (21.3-24.1) | (25.8-28.1) | (30.8-32.6) | (36.2-40.3) | (23.0-29.2) |  |
| **Probiotic Use** | 16.0<sup>a</sup> | 17.4<sup>a</sup> | 7.2<sup>b</sup> | 11.8<sup>a,b</sup> | 14.8 | NS |
| **(%)** |  |  |  |  |  |  |
| **Mean Shannon** | 4.38<sup>a</sup> | 4.35<sup>a,b</sup> | 4.27<sup>b</sup> | 4.21<sup>b</sup> | 4.35 | +0.35*** |
| **Diversity (SD)** | (0.34) | (0.31) | (0.34) | (0.36) | (0.34) |  |

## Supplementary Figures

**Supplementary Figure 1:**
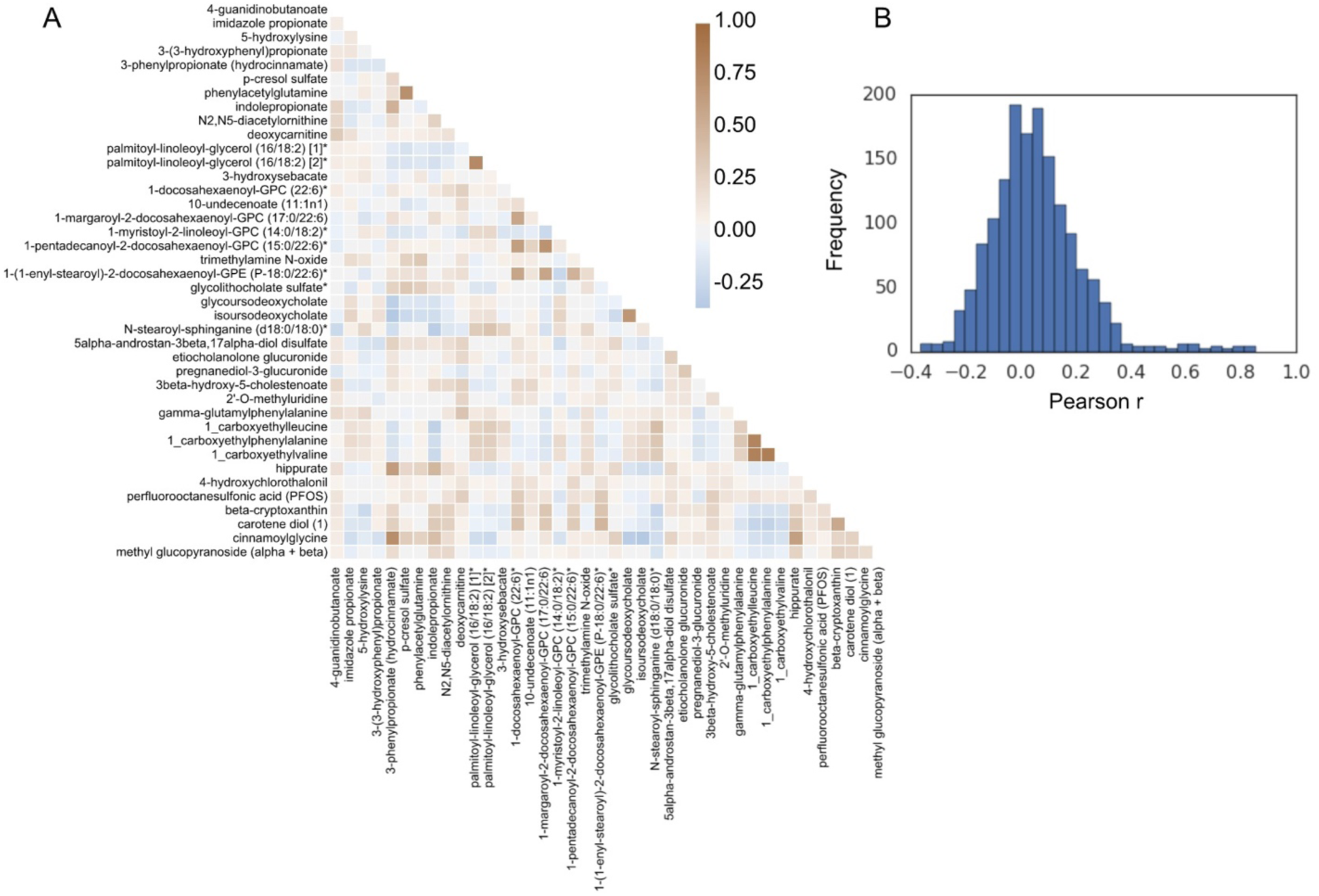
Investigating collinearity among the identified predictors of Shannon diversity in the discovery cohort. A) Heatmap showing the strength of correlation between each metabolite-metabolite pair. B) Histogram of all calculated Pearson r values for the 1560 metabolite-metabolite comparisons. Only six comparisons yielded a Pearson r value |r|>0.80.

**Supplementary Figure 2:**
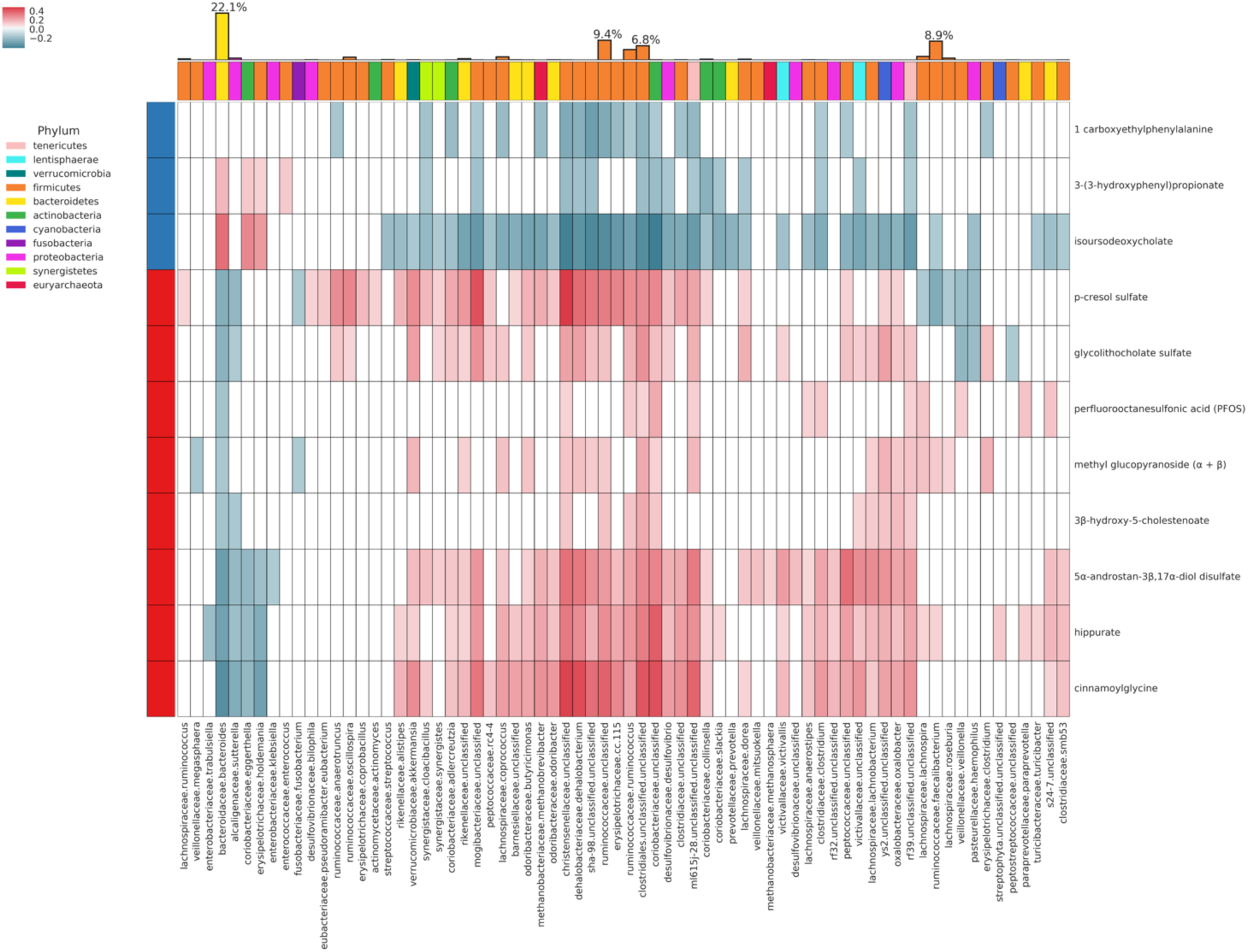
Spearman correlation of each of the 11 metabolites retained by all 10 LASSO models (rows) with microbiome genera (columns), correcting for multiple hypothesis testing (FDR<0.05). Only genera with at least one significant correlation value are displayed. Top color row labels the phylum for each genus. Left color column labels the sign of the mean β-coefficient for that metabolite across the LASSO models predicting Shannon diversity (blue - negative, red - positive). The top bar graph represents the fractional abundance of each genus, with bars colored by phylum.

**Supplementary Figure 3:**
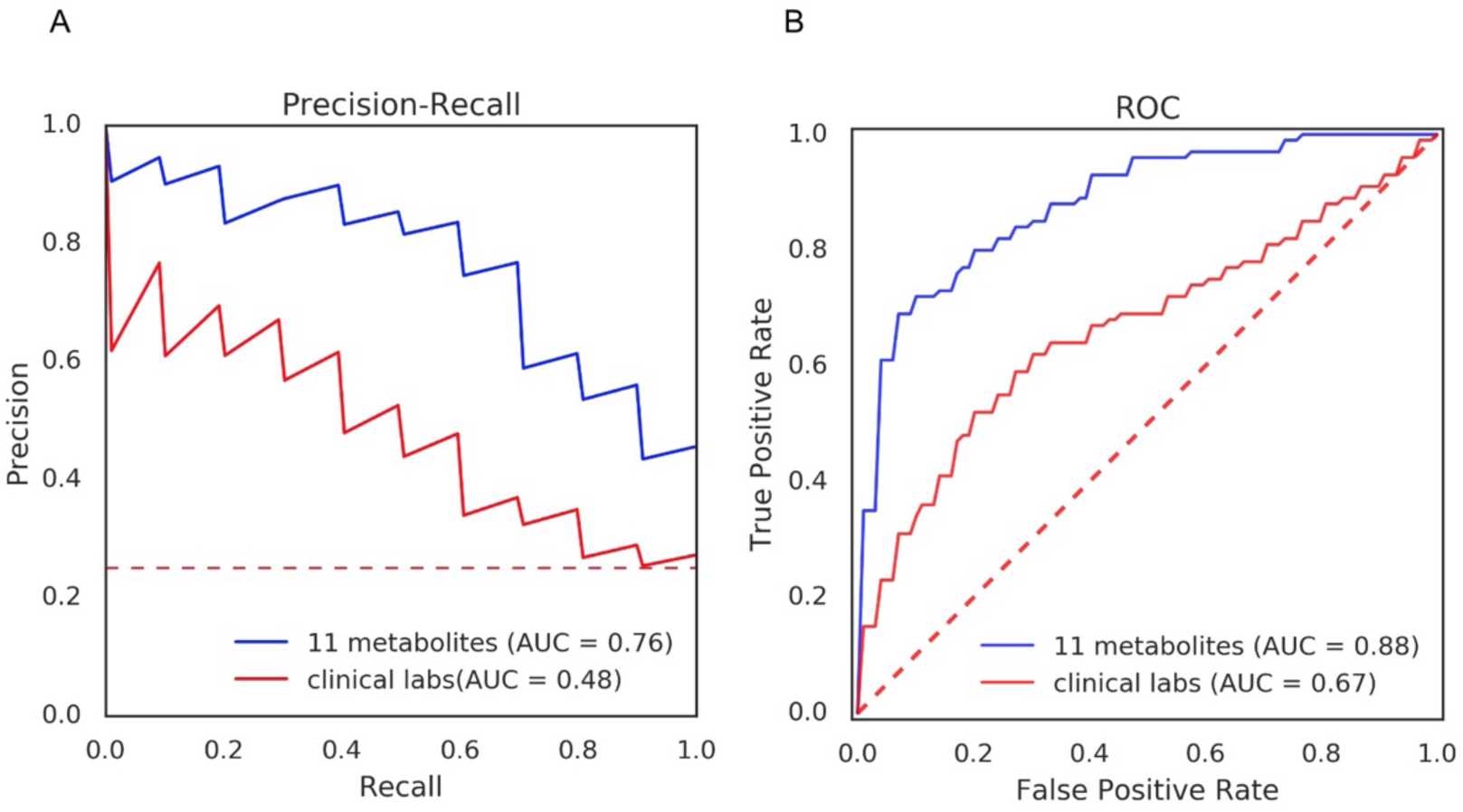
Comparison of (A) precision-recall curves and (B) receiver operator characteristic (ROC) curves for clinical labs and 11 blood metabolites classifying participants in the bottom quartile of Shannon diversity using 10-fold CV implementation of Random forests.

**Supplementary Figure 4:**
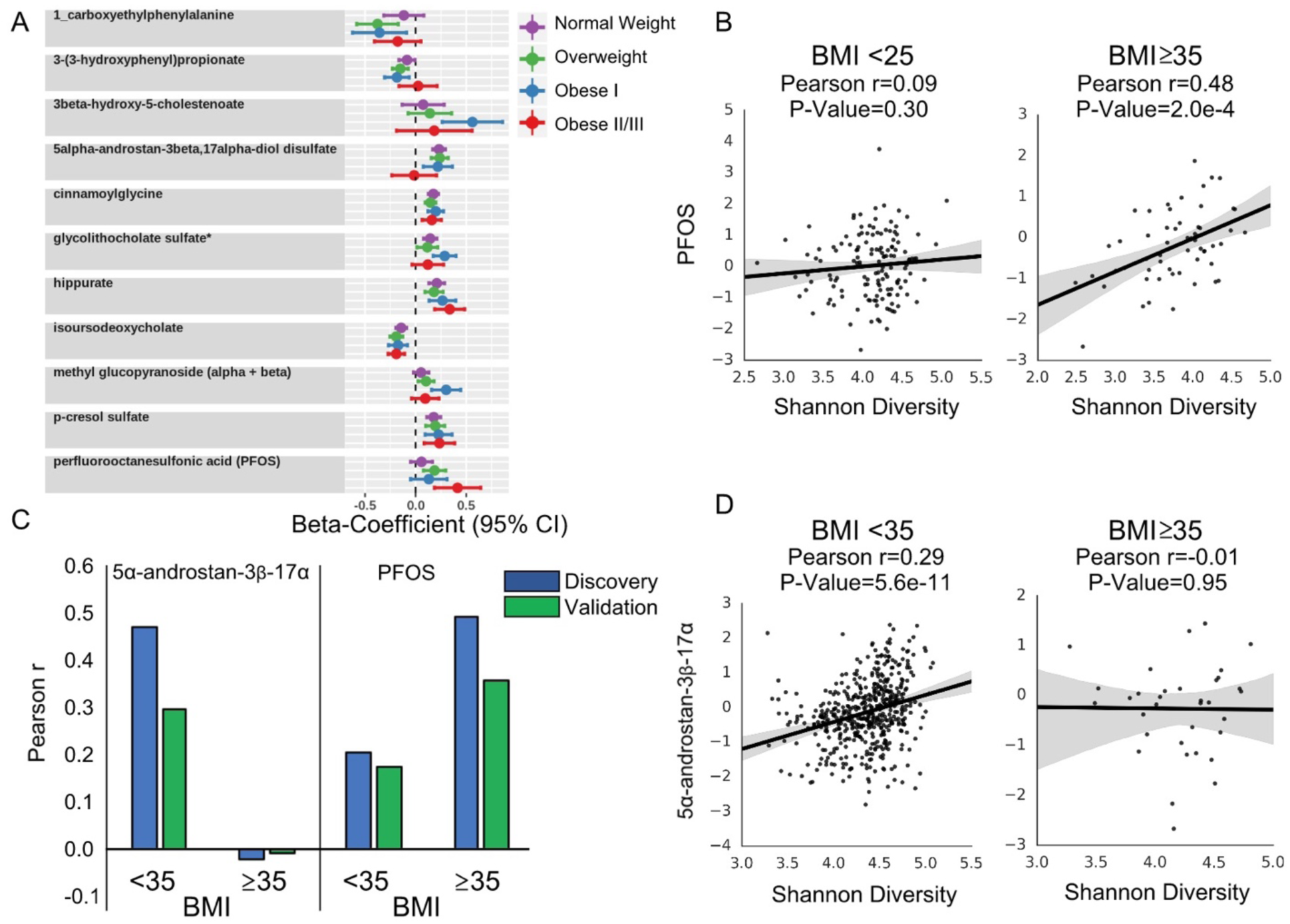
Relationship of the blood metabolome and Shannon diversity changes across BMI classes. (A) *β*-coefficients for each of the 11 metabolites retained by all 10 LASSO models from an OLS regression model with Shannon diversity as the dependent variable and sex and age included as covariates in the discovery cohort. The cohort was stratified based on defined BMI cutoffs and models were fitted independently for each BMI class. (B) Scatter plot of PFOS and Shannon diversity for participants whose BMI is less than 25 (normal weight), and greater than 35 (Obese II/III) in the discovery cohort. C) Comparison of strengths of correlations for 5α-androstan-3β-17α and PFOS with Shannon diversity across obesity in the discovery and validation cohorts. D) Scatter plot of 5α-androstan-3β-17α and Shannon diversity for participants whose BMI is less than or equal to 35, and greater than 35 (Obese II/III) in the validation cohort. Pearson r and P-values are shown. *Abbreviations: 5α-androstan-3β-17α: 5α-androstan-3β-17α-diol disulfate; PFOS: perfluorooctanosulfic acid*.

**Supplementary Figure 5:**
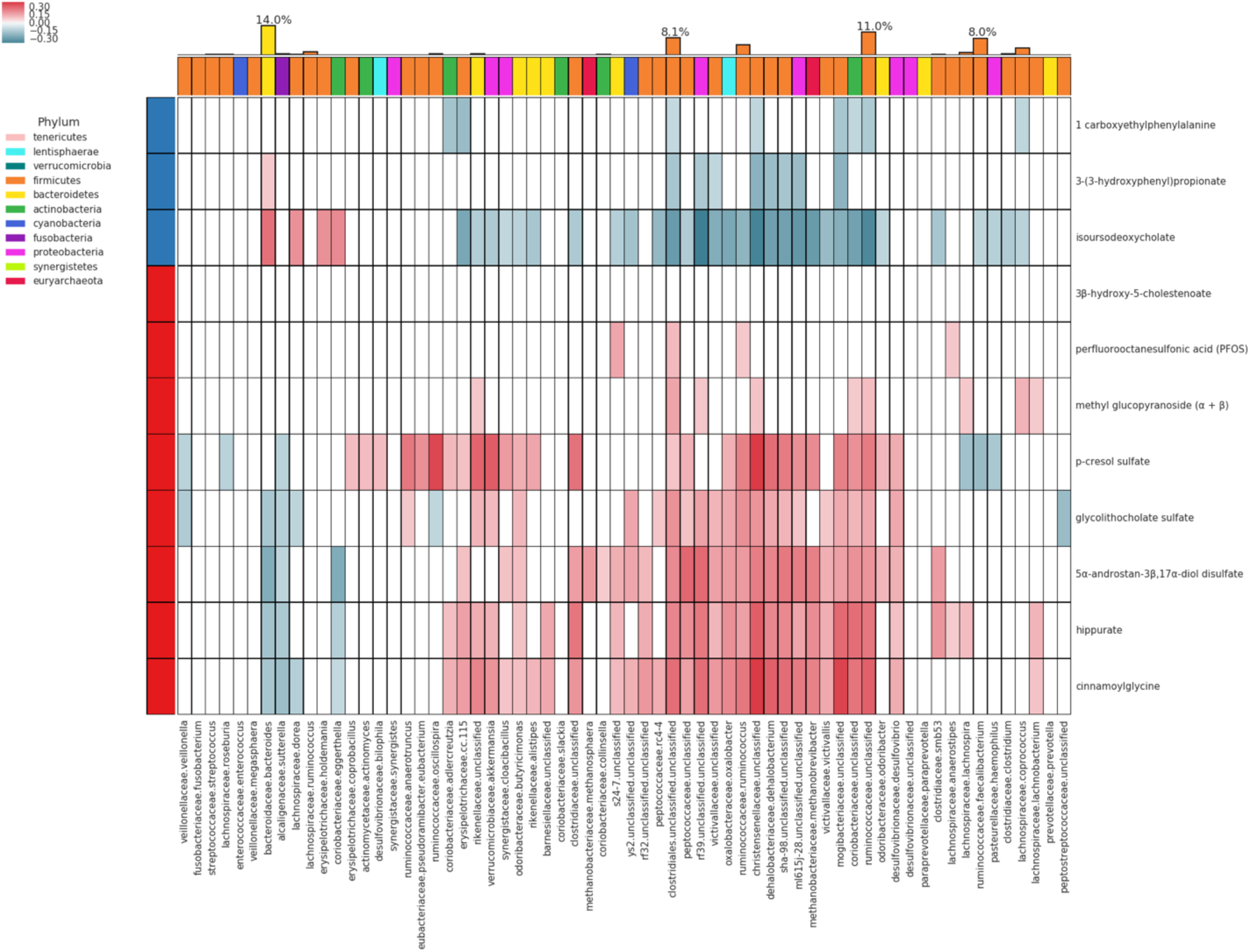
Spearman correlation of each of the 11 strongest metabolites identified in the discovery set (rows) with microbiome genera (columns) in the validation cohort, correcting for multiple hypothesis testing (FDR<0.05). Only genera metabolite correlations that were significant in the discovery cohort were considered. Top color row labels the phylum for each genus. Left color column labels the sign of the mean β-coefficient for that metabolite across the LASSO models predicting Shannon diversity (blue - negative, red - positive). The top bar graph represents the fractional abundance of each genus in the validation cohort, with bars colored by phylum.

**Supplementary Figure 6:**
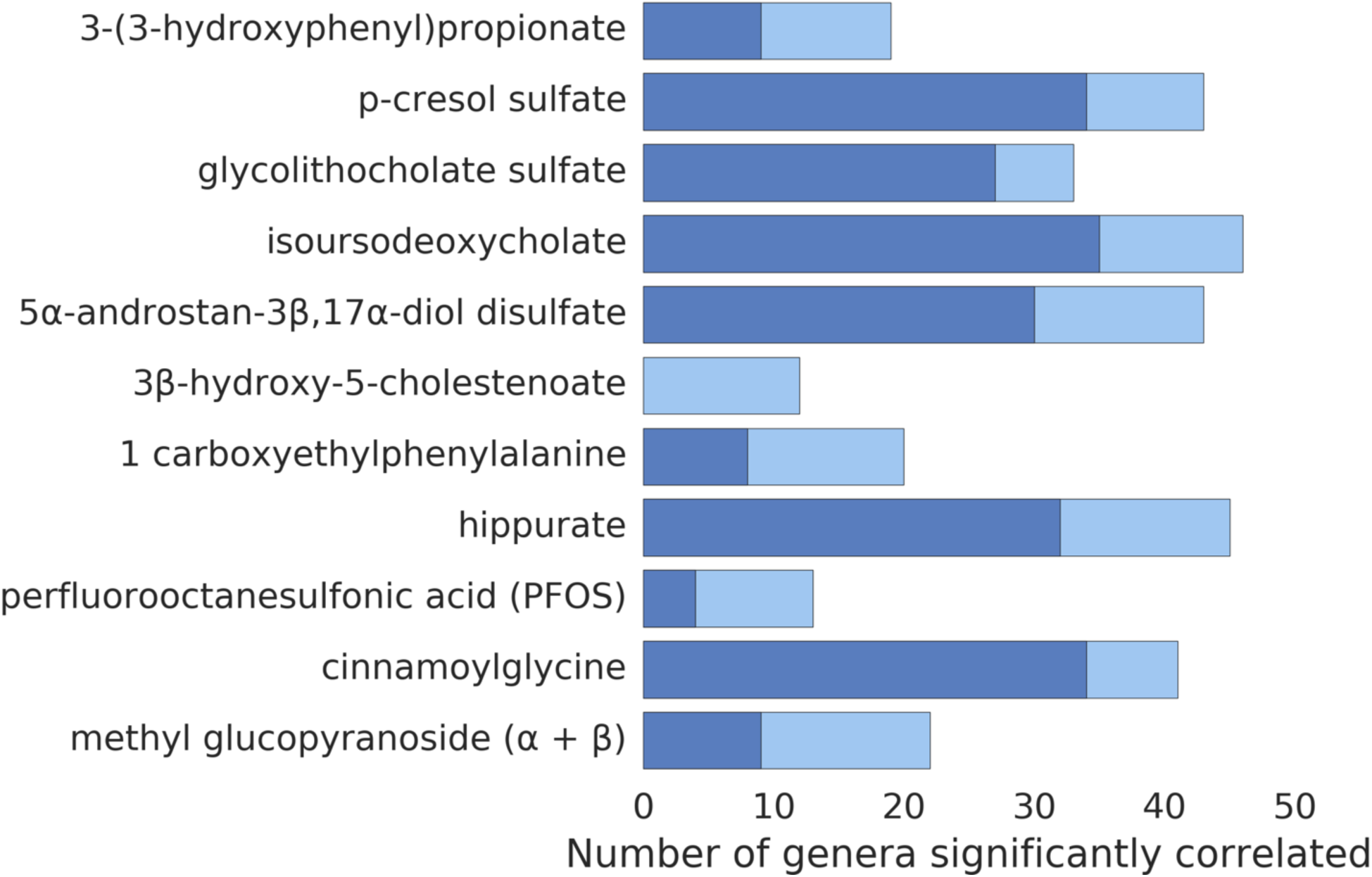
The number of significant Spearman correlations of each of the 11 metabolites retained by all 10 LASSO models (rows) with microbiome genera in the discovery (light blue) and validation (dark blue) cohort, correcting for multiple hypothesis testing (FDR<0.05). Only correlations significant in the discovery cohort were considered for validation.

